# Egyptian rousette bat humoral immunity to H9 influenza hemagglutinin

**DOI:** 10.64898/2026.06.04.730146

**Authors:** Anne A. Roffler, Bruno Bonnettaz, Emerson Glassey, Connor L. Murphy, Sophia L. Sordilla, Ceili L. Peng, Daniel P. Maurer, Eda Altan, Kristian M. Forbes, Tarja Sironen, Goran Bajic, Aaron G. Schmidt

## Abstract

In mammals, antibodies are central to antiviral defense, but they can also impose selective pressure that drives viral evolution. The interplay between viral antigenic variation and host antibody diversification constitutes a molecular arms race that influences pathogenicity, transmission, and spillover risk. Bats are reservoirs for zoonotic viruses with pandemic potential yet they appear to tolerate infection without overt disease. Although distinctive features of bat innate immunity have been described, the role of adaptive immunity—particularly antibody-mediated responses—remains largely undefined. Moreover, how antibody evolutionary pressure operates in bats is unknown, in part because tools to interrogate bat B cell responses at the monoclonal level are limited. Here, we developed a yeast surface display library of bat antibodies derived from splenic RNA of wild-caught Egyptian rousette bats to interrogate humoral responses to the bat-derived H9 influenza hemagglutinin. We isolated monoclonal antibodies recognizing the hemagglutinin (HA) antigen and defined their gene usage, somatic hypermutation frequency, binding affinities, and breadth. We then used cryo-EM to structurally characterize three bat antibodies in complex with HA engaging distinct antigenic sites. Together, these data enable direct comparison with human anti-influenza antibodies highlighting similarities in humoral immunity across mammals and provides a tool to examine bat antibody responses to other potential zoonotic viruses.

## INTRODUCTION

Bats harbor zoonotic viruses with pandemic potential, including filoviruses, influenza viruses, and coronaviruses^1^. While these viruses can cause severe disease in humans, they appear to have minimal adverse effects in their bat hosts. Understanding how bats remain largely asymptomatic and how their underlying immunological responses may differ from other mammals may inform therapeutic discovery and vaccine design. While previous studies largely focused on characterizing bat innate immune responses, less is known about the role of their adaptive immunity. A key component of the adaptative immune response include antibodies responsible for recognizing invading pathogens. In humans, antibody-mediated neutralization is an established correlate of protection. Bats similarly mount serum antibody responses following infection, and presence of virus-specific antibodies is correlated with viral clearance and reduced transmission^2–6^. Upon secondary infection, bats can generate a rapid antibody response indicative of a memory B cell compartment^6,7^.

The diverse naïve or germline antibody repertoire is subsequently diversified upon pathogen exposure. Initial diversity is established through genetic recombination of V-, D-, and J-and V- and J-genes in the heavy and light chains, respectively; the latter diversification occurs through somatic hypermutation (SHM). In humans, B cell receptors (BCR) undergo SHM and positive selection in germinal centers to improve antigen affinity and specificity^8,9^. While SHM has been suggested for several bat species, including the Jamaican fruit bat (*Artibeus jamaicensis)* and the little brown bat (*Myotis lucifugus*) they appear to undergo this process less frequently than humans. It was therefore suggested that bats elicit a lower affinity humoral response, primarily relying on V(D)J recombination for antibody repertoire diversification^10,11^. These data were based on serum analyses, genomic sequencing, and BCR sequencing. However, there is a limited understanding of the bat antibody repertoire at the monoclonal level including biochemical, biophysical, and functional analyses, in part due to the lack of necessary tools and reagents. Annotating the immunoglobulin germline loci is essential to facilitate antibody repertoire analyses, but few bats have had their immunoglobulin loci annotated or their genomes sequenced– though the Egyptian rousette bat (ERB; *Rousettus aegyptiacus*) heavy chain locus is a notable exception^12^.

ERBs have been a focus for serological surveillance and vaccination studies because of the diverse viruses detected in them including coronaviruses, filoviruses, and ortho- and paramyxoviruses ^6,13–18^. A distinct H9N2 (A/bat/Egypt/381OP/2017) influenza A virus (IAV) was isolated from ERBs, likely resulting from an avian-to-bat transmission^19,20^. Bat H9N2 IAV has since been identified ERBs across Africa, including South Africa and Kenya^21^ and there is a high seroprevalence of H9N2 IAV in these bats^22^. Despite retaining α2,3 sialic acid receptor specificity in its hemagglutinin (HA), bat H9N2 IAV transmits efficiently between ferrets, making this mammalian adapted, antigenically distinct H9N2 IAV a potential pandemic concern^23,24^. Avian H9N2 has been identified across Asia, the Middle East, and West Africa, causing sporadic human infections^25,26^. There has also been high seroprevalence to avian H9N2 in poultry workers^27^. Avian H9N2 IAV has reassorted with other circulating IAVs to cause zoonotic outbreaks^28–30^. While human and murine responses to avian H9N2 IAV, and IAVs broadly, have been well characterized after both infection and vaccination, little is known about bat antibody responses to bat H9N2 IAV and bat-to-human transmission potential.

Here, we characterized the genetic and molecular features of the ERB antibody repertoire from isolated splenocytes and constructed a yeast display library to identify antibodies specific for bat H9N2 HA. We identified 35 V_H_ genes that have not been previously described and determined cryo-EM structures of three representative bat monoclonal antibodies in complex with bat H9 HA. These latter data show similarities between bat and human antibody responses to HA and provide molecular insights into viral antigen recognition by bat humoral immunity. Furthermore, this yeast-display platform enables high-throughput isolation and molecular characterization of bat monoclonal antibodies targeting surface glycoproteins from other viruses of zoonotic potential.

## RESULTS

### Identification and annotation of Egyptian rousette bat variable heavy genes

To gain insight into the composition of the Egyptian rousette bat (ERB) transcribed antibody repertoire, we isolated and sequenced the B cell receptor (BCR) repertoire from the spleens of two wild-caught ERBs in Taita-Taveta county, Kenya. While the ERB heavy chain locus was previously annotated, the light chain loci were not^12^. We therefore used 5’ rapid amplification of cDNA ends (5’ RACE) with heavy chain constant and lambda or kappa light chain constants gene-specific primers to enrich, and subsequently sequence, antibody transcripts (**Fig. 1A**). Unlike the Jamaican fruit bat^11^, the ERB has a robust transcribed kappa light chain repertoire. Approximately 16 million paired reads were filtered to 823,981 heavy chain, 1,285,200 lambda, and 1,920,159 kappa high quality BCR sequences, nearly all of which were unique (**fig. S1A**).

**Figure 1.**
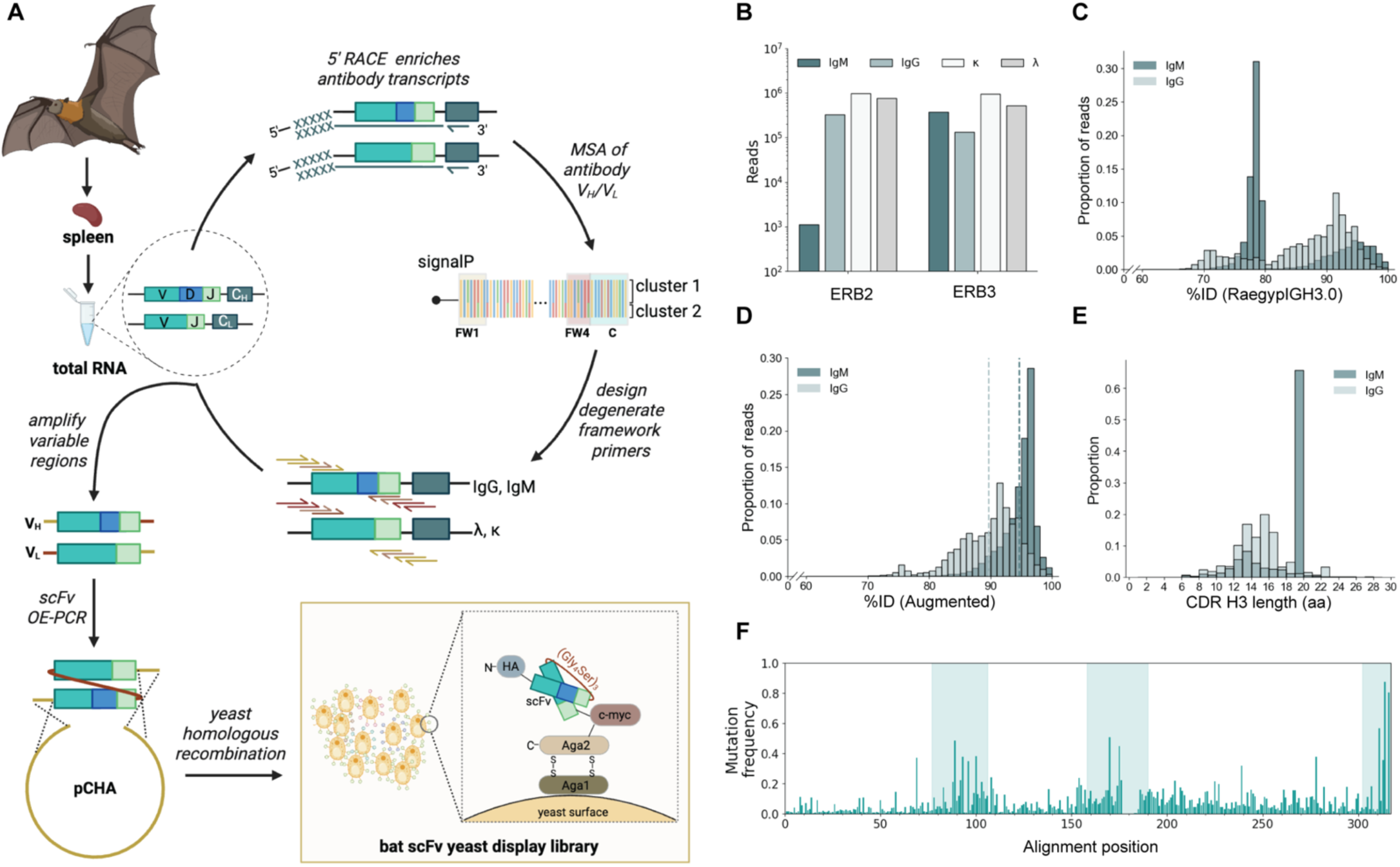
Combinatorial yeast display library expresses the Egyptian Rousette bat antibody repertoire. **(A)** Schematic for generating a naïve bat scFv yeast display library from Egyptian Rousette bat (ERB) total splenic RNA. **(B)** Paired read counts for IgM, IgG, kappa, and lambda transcripts for each ERB. **(C)** Percent nucleotide identities (%ID) of IgM and IgG transcripts to their nearest V_H_ gene as determined using the V_H_ genes previously described^12^. **(D)** Percent nucleotide identities (%ID) as determined using the V_H_ genes described here. Vertical dotted lines indicate mean %ID. Transcript-IGHV gene matching performed using IgBLAST 1.22.0. **(E)** CDR H3 length distribution as a proportion of all IgM or IgG transcripts. **(F)** SHM positional frequency along V_H_ gene alignment. CDR 1, CDR 2, and V-gene encoded start of CDR 3 highlighted.

We determined the gene identity of germline (IgM) and class-switched (IgG) BCRs. While there was comparable coverage of IgM, IgG, kappa, and lambda transcripts from both bats, ERB2 had substantially more IgG than IgM transcripts sequenced (**Fig. 1B**). The heavy chain transcripts were mapped to their closest germline relative using a custom IgBLAST database of ERB variable heavy (V_H_) genes^12^. However, there was a subset of both IgM and IgG sequences (∼41%) that shared less than 80% nucleotide identity to their nearest V_H_ gene relative (**Fig. 1C**). Because a large fraction of these sequences was from the IgM repertoire, it is unlikely due to somatic hypermutation (SHM); rather, these sequences appeared to arise from ERB V_H_ genes absent from the annotated IGH locus. We therefore queried the ERB genome (Raegyp2.0)^31^ using BLAST+^32^ and identified 35 putative V_H_ genes across 12 Raegyp2.0 scaffolds (totaling 419482 bp), 28 of which had been annotated by automated NCBI computational analysis (**table S1)**. All putative V_H_ genes clustered into V_H_ subgroups I, II, and III (**fig. S1B**)^33^. One such identified V_H_ gene, 48_VH7-like, is identical to the published V_H_7-4 gene sequence, aside from the leader sequence. All other identified putative V_H_ genes were genetically distinct from known ERB V_H_ genes (**fig. S1B**). Augmenting the IgBLAST database with these newly identified V_H_ genes reduced the proportion of V_H_ sequences with less than 80% nucleotide identity to their closest V_H_ gene match from ∼41% to ∼3% (**Fig. 1D**). However, ∼2% of total V_H_ sequences with <78% identity remained and may represent transcripts that originate from an unidentified V_H_ gene(s); sequences with <80% nucleotide identity to the augmented V_H_ gene database were excluded from downstream analyses.

### Genetic features of the Egyptian rousette bat B cell repertoire

We next assessed the genetic features of the sequenced B cell receptor transcripts. For the IgM and IgG transcripts, the mean percent identity to germline was ∼95% and ∼89%, respectively (**Fig. 1D**). The tail distribution of somatically mutated IgM sequences possibly represents antigen experienced IgM memory B cells^34–36^. A small proportion of IgG transcripts were identical to germline (<0.01%) and likely represent class-switched antigen experienced naïve B cells prior to entering the germinal center^37^. To estimate the level of SHM in each ERB, we assessed the frequency of nucleotide differences from germline V_H_ genes in IgM and IgG BCRs (**fig. S1C**). As expected, we observed less SHM in the IgM BCR repertoire than in the IgG BCR repertoire. The degree of SHM differed between the two bats, primarily in the IgG BCR repertoire. The median IgG SHM frequency is 0.08±0.037 and 0.127±0.064 for ERB2 and ERB3, respectively. For an average V_H_ length of 300 bp, this corresponds to 24 to 38 nucleotide mutations per transcript. Although rare, these levels of SHM have been observed in humans, but the majority of encoded human IgG have ∼15 nucleotide mutations^38,39^. Overall, the bat class-switched repertoire is more somatically mutated than their IgM repertoire.

In mammals, SHM primarily occurs in the complementarity-determining regions (CDRs) of the V_H_ and V_L_ genes^40,41^. To determine whether somatic mutations similarly clustered in CDRs in bats, we tabulated the differences between the sequenced transcripts and its nearest germline variable gene and plotted those differences as a function of nucleotide position (**Fig. 1F**). Consistent with humans, mutations occurred within the CDRs, with the highest mutational frequency found in CDR H3. The V_H_3-71.1, V_H_3-71.2, and 49_VH3-like genes have an additional 9 nucleotides in the CDR H2 and for all 2943 sequences mapping to these germline V genes, no mutations were found within this region (**Fig. 1F**). In the framework regions outside of the CDRs, there were three hotspots at nucleotide positions 70, 240, and 279; unlike humans, we observed consistent SHM across framework region 3^38^.

CDR H3 lengths ranged from 6 to 28 amino acids (aa) (**Fig. 1D**) for both the IgM and IgG sequences. Unlike humans, ERB CDR H3s are not noticeably longer in their IgM compared to IgG repertoires^42,43^ and are consistent with observations in the Jamaican fruit bat^11^. The IgM sequences had a large population of CDR H3s with 19aa from the V_H_4 gene family (1_VH4-like, DH2-15, JH4.3), almost all sharing an identical CDR H3 sequence. This was not observed in the IgG repertoire, which had a mean CDR H3 length ∼14aa, similar to human repertoires^39,44^ and longer than mouse repertoires^45^. For kappa light chains, most unique sequences had a 9aa long CDR L3, while the lambda light chains ranged from 8-16aa with a majority at 11aa (**fig. S1D)**. Similar with the heavy chain repertoire, the bat light chain CDR lengths are more closely related to what is observed in humans than in mice^46,47^.

### Generation of wild caught Egyptian rousette bat antibody yeast display library

To identify and characterize antigen-specific bat monoclonal antibodies (batAbs), we adapted a yeast surface display platform used previously to study naïve human antibody repertoires^48,49^. Degenerate framework 1 and framework 4 primers were designed to amplify the V_H_ and V_L_ chains from each bat (**fig. S2 and table S4**). Because we did not have proper pairing between the heavy and light chains from the sequencing, all V_H_ sequences were paired with either kappa or lambda sequences using a glycine-serine linker to generate single chain variable fragments (scFvs). V_H_-V_k_ and V_H_- V_λ_ scFvs were separately electroporated into AWY101 yeast^50^ to generate two scFv yeast display libraries (**Fig. 1A**). The V_k_- and V_λ_-containing scFv libraries expressed at ∼24.6% and ∼32.1%, respectively, we thus combined the two libraries for a total theoretical library diversity of ∼670 million scFvs with ∼28% library expression (**fig. S2**). This yeast display library is representative of the diverse transcribed antibody repertoire from both bats.

### Characterization of H9 hemagglutinin-specific bat antibodies

We used the yeast display library to identify batAbs specific to the influenza HA from the H9N2 A/bat/Egypt/381OP/2017. We designed an H9 HA soluble ectodomain with stabilizing mutations in the HA2 and an R327A mutation in the fusion peptide to impede HA proteolytic activation, resulting in recombinant expression (**fig. S3**)^51^. Using flow cytometry, batAbs were isolated by titrating the H9 HA probe from 1µM to 1nM in successive rounds of enrichment (**Fig. 2C and fig. S4A,B**) and specificity to H9 HA was ensured by using two different fluorophores (**Fig. 2D and fig. S4C**).

**Figure 2.**
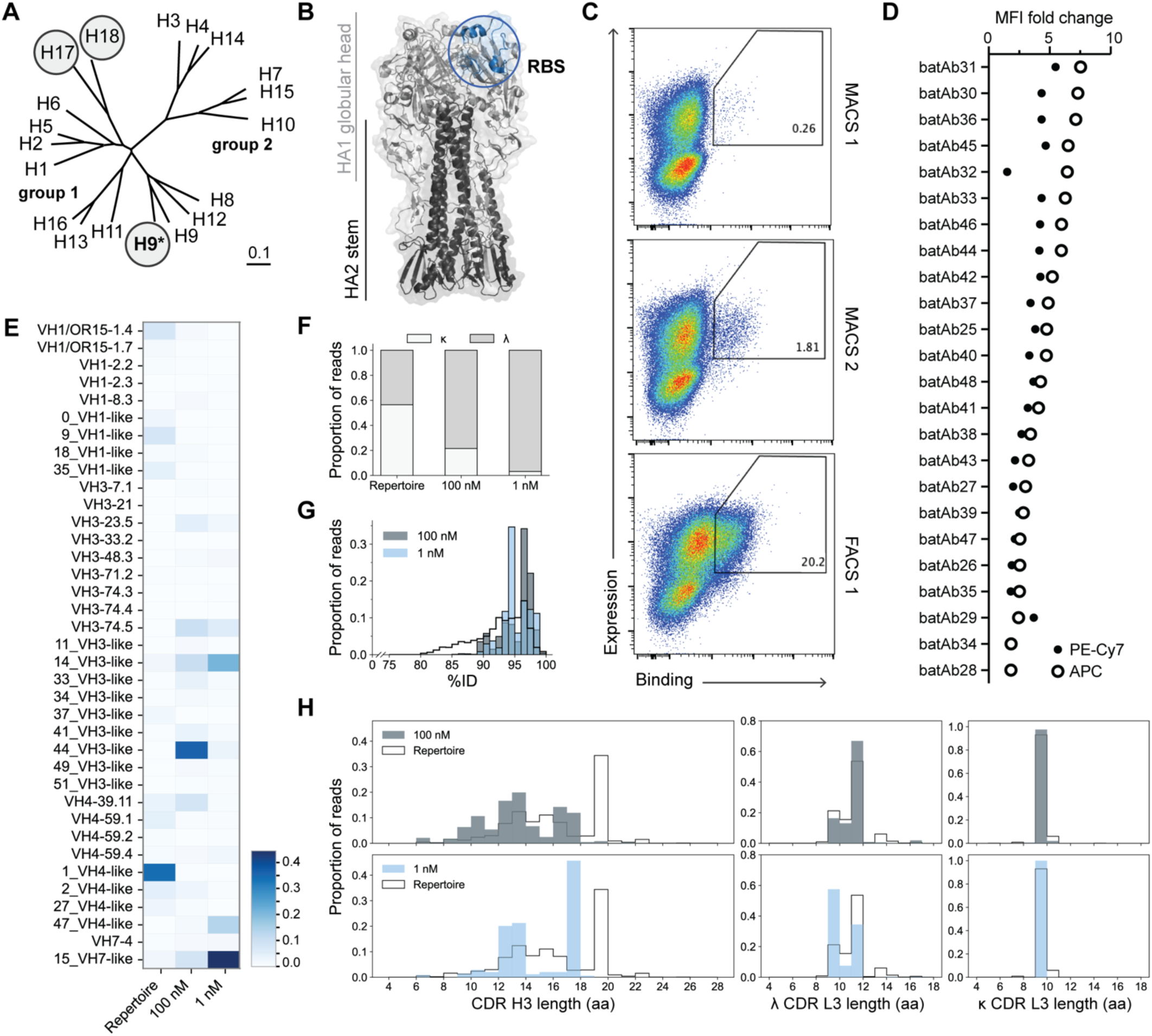
H9-sepcific bat antibody isolation and characterization. **(A)** Phylogenetic tree of group 1 and 2 influenza HAs with bat-specific subtypes circled. A/Bat/Egypt/381OP/2017 is indicated with asterisk. **(B)** H9 HA structure with head, stem, and RBS marked (PDB 1JSD). **(C)** H9 HA reactive bat antibodies were enriched over successive rounds of magnetic activated cell sorting (MACS) and fluorescence-activated cell sorting (FACS) gated on binding (H9 HA:biotin, strep-APC) and expression (c-myc, AF488); a representative plot from MACS 1 output, MACS 2 output, and FACS 1 output stained with 250 nM H9 HA shown to demonstrate round over round enrichment. **(D)** Fold change of median fluorescence intensity (MFI) of single clones gated on expression (c-myc; AF488) and binding (strep APC, strep PEcy-7). MFI calculated as fold change over MFI of c-myc; secondary-only control. **(E)** V_H_ gene usage, **(F)** light chain biasing, and (**G**) percent nucleotide identity to germline V_H_ genes of splenic antibody repertoire, moderate affinity (100nM) and high affinity (1nM) H9 HA reactive batAb populations as shown as proportion of total unique sequences for each population. **(H)** Heavy and light chain CDR amino acid (aa) length distribution of moderate affinity (100nM, top row) and high affinity (1nM, bottom row) H9 HA reactive batAbs compared to splenic antibody repertoire CDR length distribution, represented as proportion of total unique sequences for each population.

To assess the genetic background of the H9 HA reactive bat antibody repertoire, we deep sequenced the output of our terminal round of sorting against 100nM H9 HA and obtained 1153 unique batAb sequences. We did an additional enrichment campaign using H9 HA at 1nM to isolate higher affinity clones (**fig. S4**). From this additional sort, we obtained sequences for 375 unique batAbs, 217 of which overlapped with the initial 100nM-reactive population. We removed overlapping sequences and analyzed the 100nM “moderate affinity” population and 1nM “high affinity” population independently. To further ensure that the analyzed sequences are H9 HA-specific, at each round of sorting, we also sorted and sequenced nonspecific yeast clones and removed them from subsequent analyses.

The moderate affinity population predominantly arose from three putative V_H_ genes: 44_VH3-like, 14_VH3-like, and the previously published V_H_3-74.5. For the two putative V_H_ genes, the closest published V_H_ relatives are V_H_3-21 (96.9% identity) and V_H_3-71.1 (70.3% identity), respectively (**Fig. 2E and fig. S1**). Antibodies that used V_H_ 14_VH3-like, 15_VH7-like, and 47_VH4-like were selectively enriched in the high affinity population (**Fig. 2E**). These latter two putative V_H_ genes are closely related to V_H_7-4 (99.7% identity) and V_H_4-59.1 (99% identity), respectively, and encoded most antibodies enriched for affinity during our sorting campaign. Although our splenic BCR repertoire analysis from both bats showed comparable usage of kappa-and lambda-light chains (∼57% kappa, ∼43% lambda), the H9-reactive antibody population selectively used lambda-light chains (**Fig. 2F**). Of the moderate affinity population, ∼78% (734/936) of unique clones were lambda-containing, which increased to ∼97% (363/375) in the high affinity population. Because we are unable to determine whether the enriched heavy chains were IgG or IgM, we first determined the median mutation frequency of all heavy chain sequences from both bats to be 0.067±0.04. (**Fig. 2G**). This was greater than both the isolated high and moderate affinity populations of H9 HA which was 0.054±0.021 and 0.034±0.025, respectively (**Fig. 2G**). These data suggest that increased affinity to H9 HA is associated with more SHM but that overall, H9 HA reactivity appears to require minimal SHM for affinity and specificity (**Fig. 2G**).

We next determined the heavy and light chain CDR3 lengths of the H9 HA-specific batAbs and compared them to the splenic BCR repertoire from both bats. We found that high affinity H9 HA reactive batAbs had CDR H3 lengths of 12, 13, and 17aa (**Fig. 2H**). Notably, 17aa CDR H3 heavy chains comprised a small proportion of the bat splenic BCR repertoire but became selectively enriched in the moderate and high affinity H9 HA-reactive populations. Lambda H9 HA-specific batAbs had predominantly 9 and 11aa CDR L3s, with high affinity lambda light chains predominantly using a 9aa CDR L3 (**Fig. 2H**). The kappa light chain H9 HA reactive batAbs had 9aa CDR L3s, as was also observed on the repertoire level (**Fig. 2H**). Without an annotated ERB kappa locus, we used IgBLAST to analyze the kappa sequences against the human Ig database and found representation from four out of five human kappa J genes, with IGKJ2 accounting for ∼53% of the sequences. These data are consistent with CDR L3 length restriction observed in the human kappa repertoire^52^. Overall, we found that the H9 HA-specific subset of the total bat antibody repertoire were almost entirely lambda light antibodies paired with diverse V_H_ genes and were somatically mutated.

### Recombinant bat antibodies bind H9 HA with high affinity

To obtain binding affinities, we recombinantly expressed 24 bat IgGs downselected to reflect the polyclonal H9 HA-specific repertoire with representation from diverse variable region gene segments and unique heavy chain-light chain pairings. Only 7 out of 24 bound H9 HA with a median effective concentration (EC_50_) ranging from ∼1 nM to ∼119 nM and a mean of ∼35 nM (**Fig. 3A and fig. S5**). This was rather surprising given that the corresponding single yeast clones of the batAbs bound to H9 HA with high affinity (**fig. S4C**). This discrepancy is likely attributed to the highly avid nature of the yeast display platform. We further defined the epitope using an H9 HA head-only construct and found that batAb55, 31, and 58 bound with comparable affinity to both H9 HA head and full-length constructs suggesting their epitope(s) are in the HA head (**Fig. 3A**). No recombinant batAb reacted nonspecifically with streptavidin **(fig. S5**).

**Figure 3.**
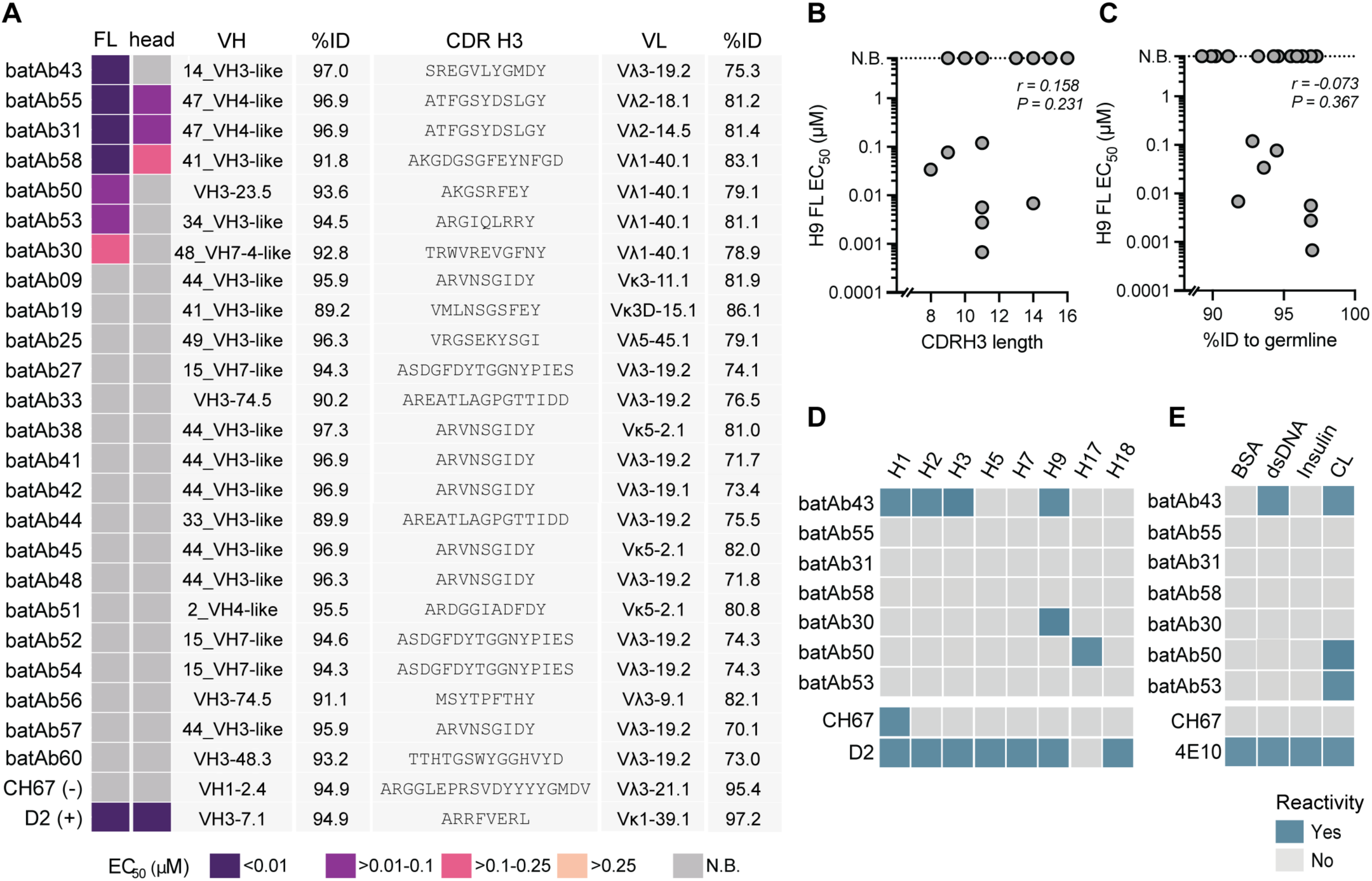
Features of H9 HA-reactive bat antibodies. **(A)** ELISA EC_50_ (nM) binding heat map of 24 batAbs expressed as bat IgG1 to H9 HA full length ectodomain (FL) and head domain (head). N.B., no binding. FL- and positive head-based ELISAs were performed in technical triplicate. For H9 HA FL nonreactive batAbs, negative binding to H9 head confirmed in singlicate. Closest ERB V_H_ and human V_L_ gene shown, (human antibodies CH67 and D2 used as controls are relative to the human V_H_). %ID., percent nucleotide identity. **(B)** Pearson correlation analysis of H9 HA FL affinities and CDR H3 length (IMGT) (n=24). **(C)** Pearson correlation analysis of H9 HA FL affinities and percent identity to germline. %ID., percent nucleotide identity (n=24). **(D)** Cross reactive ELISAs performed at a single mAb concentration (50μg/mL) in triplicate against HAs. Reactivity was defined by an OD_450_≥ 0.40. **(C)** Polyreactive ELISAs performed at a single mAb concentration (50μg/mL) in triplicate; polyreactivity was defined by an OD_450_ ≥ 0.40. CL., cardiolipin.

BatAb31 and batAb55 share heavy and light chain genes and differ by only 6 amino acids in the light chain. We also enriched for a population of batAbs that use 44_VH3-like V_H_ gene, share the same CDR H3, but are paired with unique light chains (Vλ3-19.2, Vκ5-2.1, or Vκ3-11.1) (**Fig. 3A**). This likely suggests a heavy chain CDR H3-mediated interaction with H9 HA that is either light chain independent or potentially multiple binding approaches to H9 HA.

Without annotated light chain loci available to us, we used IgBLAST to analyze the batAb light chain sequences against the human light chain database and found that ERB lambda and kappa V_L_ genes are highly divergent from human genes, ranging from ∼70-86% identity (**Fig. 3A**). We next compared our lambda light chain sequences against a custom IgBLAST database of Jamaican fruit bat V_L_ genes^11^ and found a similar degree of divergence, ranging from ∼73.9-83.9% identity. This suggests divergent evolution of antibody germline genes between bat families. Of the binding population, there appeared to be no predisposition for CDR H3 length, and all paired with lambda light chains except batAb9, 19, 38, 45 and 51 (**Fig. 3B**). These batAbs ranged from ∼91.8-98% nucleotide identity to its nearest germline V_H_ gene, and in our small sample size, EC_50_ was not strongly correlated with degree of somatic hypermutation (**Fig. 3C**).

We next assessed batAbs breadth by selecting representative group 1 and group 2 influenza HAs to maximize antigenic diversity (**Fig. 3D)**. Notably, bat H9 HA has diverged significantly from other influenza subtypes, sharing only ∼72% amino acid sequence homology with avian H9 HA and ∼48% homology with the other previously identified bat-specific H17 HA. batAb43 broadly reacted against group 1 (H1, H2, and avian H9) and group 2 (H3) HAs. batAb30 was similarly cross reactive with avian H9 HA. batAb50 cross-reacted with another divergent bat-specific influenza HA (H17). All batAbs were also tested for polyreactivity using bovine serum albumin (BSA) and three common autoantigens, double-stranded DNA (dsDNA), human insulin, and cardiolipin by ELISA (**Fig. 3E**). Although batAb43 reacted with dsDNA and cardiolipin, they were not broadly cross reactive to all tested influenza HAs (**Fig. 3D**). These data show an HA specificity of isolated batAbs in a non-polyreactive manner.

### Structural basis for H9 HA recognition by bat antibodies

We determined cryo-EM structures of batFab30, batFab31 and batFab43, selected as representatives from each of the three V_H_ families, in complex with H9 HA from A/bat/Egypt/381OP/2017 to understand how these batAbs engage HA (**Fig. 4; table S3**). These batAbs were selected as representatives from each of the three V_H_ families. These structures showed that the selected batAbs recognize sites engaged by human antibodies^53–57^. Specifically, batFab30 and batFab31 engage canonical antigenic sites of the HA on the stem and head, respectively (**Fig. 4A, D)**, whereas batFab43 targets a distinct, non-canonical site at the extreme HA C-terminus (**Fig. 4G**).

**Figure 4.**
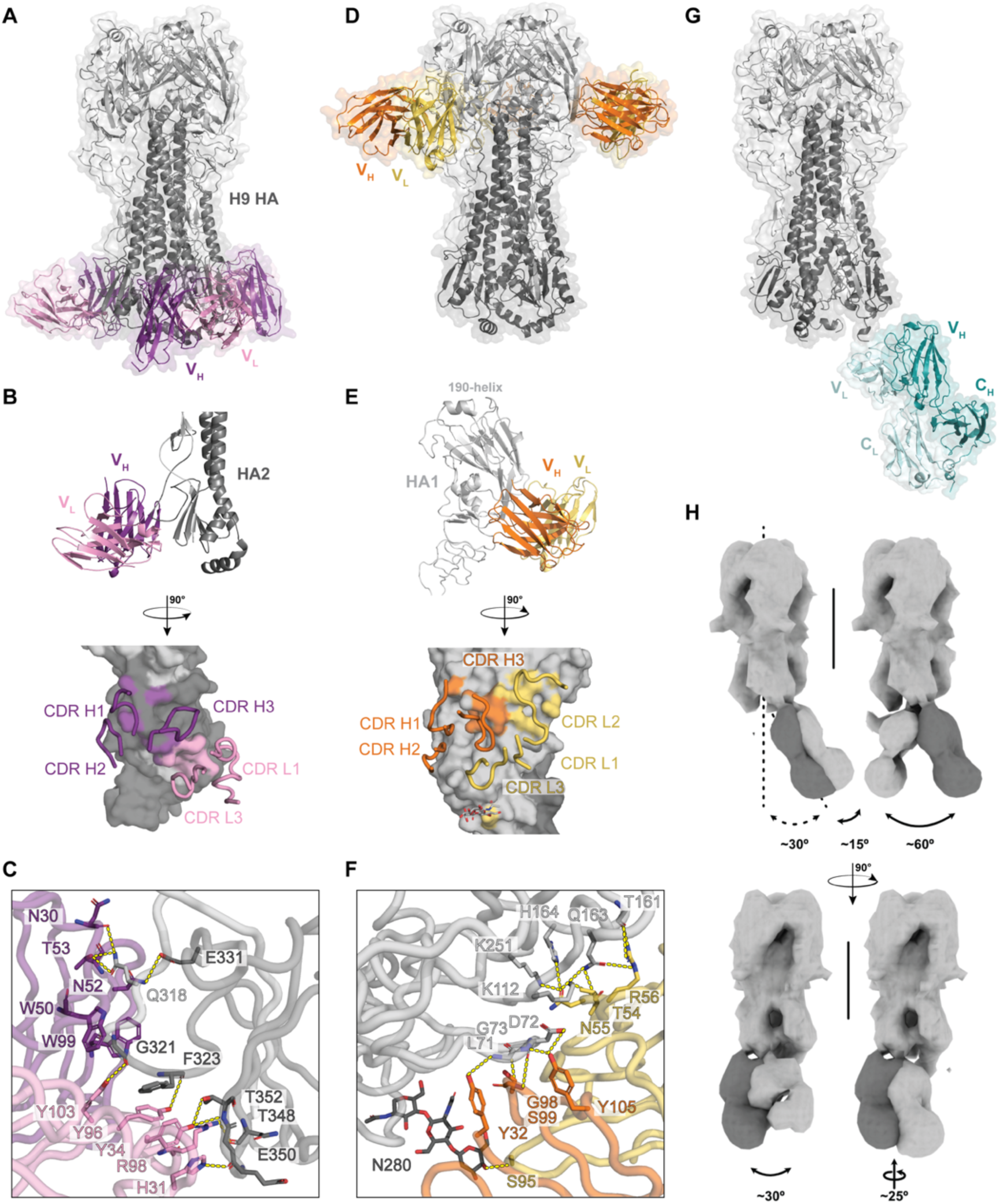
Cryo-EM structures show distinct modes of recognition by bat antibodies. **(A)** Model of batAb30 in complex with A/Bat/Egypt/381OP/2017 HA represented as ribbon and transparent surface at 2.35 Å resolution. **(B)** Fab footprint on H9 HA and CDR loops involved in binding. **(C)** Detailed batFab30:HA interactions, with key contact residues labeled and hydrogen bonds shown as dashed yellow lines. **(D)** Model of batFab31 in complex with A/Bat/Egypt/381OP/2017 HA represented as ribbon and transparent surface. 2.90 Å resolution. **(E)** Fab footprint on H9 HA and CDR loops involved in binding. **(F)** Detailed batFab31:HA interactions, with key contact residues labeled and hydrogen bonds shown as dashed yellow lines. **(G)** Model of batFab43 in complex with A/Bat/Egypt/381OP/2017 HA represented as ribbon and transparent surface at 4.5 Å resolution. **(H)** Low-resolution map corresponding to discrete orientations of batFab43. In dark grey the low-resolution map of **(G)** as reference. Dashed arrow represents angle of batFab43 main plan relative to HA three-fold symmetry axis, plain arrows represent angles between Fab main plan and representative darker Fab.

The high-resolution cryo-EM map of the batFab30-H9 HA complex showed two binding populations: a predominant 1:1 Fab:HA stoichiometry (**Fig. 4A**) and a minor 2:3 Fab:HA (**fig. S9**). BatFab 30 primarily engages the HA2 stem domain, with contributions from both heavy and light-chain CDRs (**Fig. 4B**). The interface is defined by two major polar interaction networks flanking the uncleaved fusion peptide, each contributing comparably to the binding footprint. Notably, all light-chain contacts (H31, Y34, Y96 and Y103) involve HA2 main-chain carbonyls, suggesting a structural basis for broad reactivity across H9 variants. Consistent with this observation, superposition with H9 HA from A/swine/Hong-Kong/9/98 (PDB 1JSD) that has a cleaved fusion peptide indicates that most light-chain interactions at the 360-loop are preserved upon activation (**Fig. 4C and fig. S6**). Additionally, the uncleaved fusion peptide is stabilized within the Fab binding groove by clamping of the A335 and G336 backbone by W50 and W99 from CRD H2 and CDR H3, respectively.

BatFab31 binds a canonical site on the HA head, which overlaps, in part, with the vestigial esterase domain of the HA1 subunit (**Fig. 4D**). Binding is mediated by both heavy and light chains, which contribute almost equally to an extensive polar interaction network. CDRH1 and CDR H3 interact with the HA epitope through a dense hydrogen-bonding network, primarily involving main-chain carbonyls of residue L71, D72 and G73 (**Fig. 4E, F**). On the beta-sheet domain of the HA head, CDR L2 contacts three strands and overlaps with the H9-B antigenic site described in avian H9 subtypes^58,59^. Interestingly, batFab31 is positioned over the HA N280 glycan and forms a direct contact via CDR L3 S95 (**Fig. 4E, F**). Although this interaction is observed across all 3 protomers, the partial occupancy of the glycan observed in our cryo-EM map suggests positional flexibility and a transient interaction, indicating that the glycan is not essential for binding.

In contrast, batFab43 binds a non-canonical epitope on HA. The 2D class averages (**fig. S11**) show a well-defined HA ectodomain with batFab43 flexibly attached near the stem. Although the ectodomain was resolved at an overall resolution of 2.74 Å, with uniformly high map quality across the head and stem regions, the membrane-proximal region contacted by the Fab showed lower local resolution and smeared density, consistent with substantial conformational heterogeneity. While the initial analysis suggested a continuum of poses, extensive particle classification and masking strategies^60,61^ (**fig. S11**) enabled us to reconstruct multiple maps corresponding to discrete Fab orientations at moderate resolution (**Fig. 4H, fig. S7**). BatFab43 approaches the HA ectodomain at an angle of ∼20-45° relative to the HA three-fold symmetry axis. Due to limited local resolution and noise in the Fab subvolume region, only half of the C-terminal helix of the HA could be reliably traced. Nevertheless, the data indicate that the batFab43 binds exclusively to the C-terminal helix of HA (residue 488 onward, HA0 numbering), a region that remains structurally flexible (**fig. S7**). Although 2:3 Fab:HA stoichiometry is observed in some subclasses, the predominant stoichiometry observed in the dataset is 1:3, consistent with the high mobility of the Fab and steric constrains associated with its binding pose.

As no other structures of bat antibodies are available, we compared the footprint of batFab30, batFab31 and batFab43 with previously described human and murine antibodies. BatFab30 had substantial overlap with human antibodies ADI-85666, CR8020, and MEDI8852^62–64^ and batFab31 had substantial overlap with the human antibody HD16_D04 and mouse antibodies H5M9 and 100F4^56,65–67^ (**fig. S8A, B**), indicating convergence toward defined sites of vulnerability on HA. In contrast, the angle of approach and high conformational flexibility of batFab43 closely resemble those reported for the mouse antibody 5E10^55^ (**fig. S8C, D**), suggesting a shared mode of recognition. Together, these analyses show that batAbs converge on conserved antigenic sites targeted across mammalian species, while also expanding the landscape of recognition through engagement of flexible, non-canonical regions of HA.

## DISCUSSION

Although bats can generate virus-specific humoral responses upon infection or vaccination, it remains unknown how elicited antibodies might recognize a virus. Understanding the biochemical and biophysical features of bat antibodies targeting a viral glycoprotein necessarily relies on recombinant monoclonal bat antibodies with defined antigen specificity. Here, we developed a yeast display library of bat antibodies from two wild-caught ERBs and characterized the antibody response to bat influenza H9 HA. Notably, we found structural convergence of bat antibodies to conserved regions on HA that overlap with other vaccine- and infection-elicited human antibodies.

The genomic features of the ERB immunoglobulin heavy chain locus previously described, including annotated V-, D-, and J-genes and C_H_ sequences, facilitated the work described in this study^12^. Targeted sequencing of the splenic antibody transcripts uncovered insights into expressed naïve and class-switched ERB antibody repertoires. We antibody transcripts that likely did not arise from the germline genes previously annotated in the ERB IGH locus. By comparing the transcribed repertoire to the ERB genome, we identified 35 putative V_H_ genes outside of the annotated IGH locus, with likely more V_H_ genes remaining to be annotated based on the IgG sequences still mapping with less than 80% identity to our augmented V_H_ gene database. There is evidence that multiple species within *Vespertilionida*, including the big brown bat (*Eptesicus fuscus*), have two genetically distinct and functional IGH loci on separate chromosomes^68^. Without full, contiguous sequencing of the ERB genome, it is unknown where these putative V_H_ genes localize, and whether the bimodal distribution of sequence identity in the IgG repertoire indicate a biological, highly somatically mutated class switched compartment, or is the result of an incomplete V_H_ gene database. By comparing genomic sequences with the transcribed repertoire, we described features previously unknown, including subtype-specific CDR length distributions and the frequency and location of SHM across V_H_ genes. Furthermore, we report a highly diverse light chain repertoire comprising both lambda and kappa light chains that seemed to be skewed to lambda-containing antigen-specific repertoire.

While sequencing the transcribed antibody repertoires of wild-caught or experimentally infected bats provide insight into the overall antibody compartment architecture and how it may change upon infection, it does not describe the molecular details of the antigen-specific repertoire. Identifying antibodies elicited against a viral pathogen builds a basis towards understanding how host humoral immunity recognizes and potentially clears infection as well as inhibits viral transmission. To date, there have been no bulk sequencing analyses of antigen-specific bat antibody responses. We enriched for bat antibodies specific to the H9 HA and coupled with deep sequencing of the outputs of our enrichment campaigns, have identified antibodies that either are present or could be potentially elicited upon infection; this allows a direct comparison to human responses after influenza virus infection or vaccination.

In humans, antibody responses targeting conserved viral sites can be highly convergent, resulting in antibodies with shared genetic and structural features. One such antigenic site is the influenza HA stem. Human antibodies targeting this epitope predominantly use the V_H_1-69 heavy chain, which encodes two hydrophobic residues at the tip of the CDR H2 and stabilizes binding within a hydrophobic pocket on the HA stem^69–71^. Human antibodies targeting other antigenic sites on HA have also been identified, including a broad class of genetically diverse antibodies targeting the HA trimer interface^72^, the receptor binding site (RBS)^73^, lateral patch^74^, vestigial esterase domain^67^, and anchor epitopes^75^. In ERBs we found that H9 HA-reactive bat antibodies arise from diverse V_H_ gene families, with biased V-gene usage closely related to human V_H_3-23, V_H_7-4, along with bat V_H_4-59 and putative V_H_ gene (14_VH3-like), which shares only ∼74% and ∼76% identity with its closest human V gene relatives V_H_4-4 and V_H_3-30, respectively. Though 14-VH3-like bat V_H_ gene is divergent from human V_H_3-30, a class of stem-directed human antibodies (*e.g.,* FI6) arise from V_H_3-30 lineage^76–78^ and may indicate convergent evolution towards a shared antigenic site on HA. We provide evidence that putative V_H_ genes outside of the annotated IGH locus may give rise to functional antibodies.

From the HA-batAb structures determined by cryo-EM, we found two antigenic sites commonly engaged by human and mouse antibody responses. Despite divergent gene usages, human antibody ADI-85666 and batAb30 engage an overlapping region on HA2 with common modes of recognition^62^. ADI-85666 uses V_H_4-61 and V_K_1-17 and stabilizes the fusion peptide using Y35 and R99 in its CDRH2 and CDR H3, respectively. BatAb30, however, uses V_H_7-4 and V_λ_1-40-like and stabilizes the fusion peptide using W50 and W99 from CDR H2 and CDR H3, respectively. BatAb31 footprint overlaps with both the “upper” and “lower” sites of vulnerability on the vestigial esterase domain defined by 100F4 and H5M9 mAbs, respectively, as well as HD16_D04^56,79^. The cross reactivity of batAb30 with avian H9 HA suggests that this antigenic region is conserved between avian and mammalian-adapted H9N2. We separately identified a non-canonical, hyperflexible recognition domain on H9 HA. BatAb43 uses a uniquely divergent bat V_H_ gene, sharing only ∼76% nucleotide identity with its closest human V gene relative V_H_3-30. It is unclear whether this C-terminal epitope is accessible on the virion surface. Although proximal to the N-terminal, surface-accessible loop targeted by 5E10^55^ and FISW84^80^, the HA C-terminus leads directly into the transmembrane domain that anchors the HA onto the virion surface and is further occluded than the N-terminal loop. However, with membrane-bound HA rotating up to 52° on its threefold axis^53^, it remains to be seen whether this C-terminal epitope can be accessed on the H9N2 virion surface.

Although we identified and characterized the biochemical and structural features of recombinant bat monoclonal antibodies, there are several limitations to our study. First, the limited number of wild-caught ERBs analyzed reduces the robustness and power analyses regarding the observed antibody repertoire. Second, we do not know the immune histories of these bats, which directly influences the presence and abundance of affinity-matured antibodies and thus limits conclusions about repertoire gene usage and frequency. Third, without annotated light chain loci, all V_L_ analyses performed relied on comparison to human genes. Fourth, we were unable to isolate antigen-specific bat naïve and memory B cells and used potentially non-natively paired heavy chains and light chains in our yeast display library. However, all three antibody:HA structures determined here show critical contacts mediated by both the heavy and light chains suggesting that there is likely proper-pairing within the library. Nevertheless, these data highlight features of convergent evolution between human, murine, and bat antibody responses towards influenza HA, offering a basis for understanding bat antibody-mediated recognition of a bat IAV. This study bridges previous genetic sequencing of ERB germline genes with transcribed repertoire sequencing and biochemical characterization of diverse H9-HA specific bat monoclonal antibodies, providing a functional description of the wild-caught ERB antibody repertoire. This work has broader implications for understanding bat adaptive immunity to potentially zoonotic viruses of pandemic concern.

## Acknowledgments

We thank Paul Webala, Jospeh Ogola, and Ilkka Kivistö for their assistance with bat trapping. We acknowledge support from the following sources.

P01 AI089618 (A.G.S.).

Irma T. Hirschl/Monique Weill-Caulier Trust (G.B.).

Finnish Cultural Foundation and Jenny and Antti Wihuri Foundation (T.S., K.M.F). Research Council of Finland grants no. 339510 and 358323 (T.S.).

This material is based upon work supported by the National Science Foundation Graduate Research Fellowship Program under Grant No. DGE 2140743 (A.A.R.).

Some of this research used the Glacios 2 Cryo-TEM located at the Cryo-EM CoRE facility of the Icahn School of Medicine at Mount Sinai, NIH under grant number 1S10OD036213-01.

Some of this research was performed at the National Center for CryoEM Access and Training (NCCAT) and the Simons Electron Microscopy Center located at the New York Structural Biology Center, supported by the NIH Common Fund Transformative High Resolution Cryo-Electron Microscopy program (U24 GM129539), The Simons Foundation (SF349247) and NY State Assembly.

This research was supported in part through the computational and data resources and staff expertise provided by Scientific Computing and Data at the Icahn School of Medicine at Mount Sinai and supported by the Clinical and Translational Science Awards (CTSA) grant UL1TR004419 from the National Center for Advancing Translational Sciences. Research reported in this publication was also supported by the Office of Research Infrastructure of the National Institutes of Health under award number S10OD026880 and S10OD030463.

This research has been funded in whole or part with federal funds under a contract from the National Institute of Allergy and Infectious Diseases, NIH contract 75N93019C00050 (A.G.S.). The content is solely the responsibility of the authors and does not necessarily represent the official views of the National Institutes of Health.

## Author contributions

Conceptualization: AAR, AGS, DPM

Methodology: AAR, EG, CLM, DPM

Investigation: AAR, BB, EG, CLM, EA, SLS, CLP

Visualization: AAR, BB, CLM

Funding acquisition: KMF, TS, GB, AGS

Project administration: KMF, TS, GB, AGS

Supervision: KMF, TS, GB, AGS

Writing – original draft: AAR, BB, AGS

Writing – review & editing: AAR, BB, EG, CLM, EA, SLS, CLP, AAR, KMF, TS, GB, AGS

## Data and materials availability

Wild-caught ERB antibody repertoire datasets are available at the NCBI Short Read Archive (BioProject: PRJNA1471324). The 100nM and 1nM H9-HA reactive bat scFv yeast datasets are available at BioSample: SAMN60509428-60509431. V_H_ and V_L_ sequences for isolated mAbs are available at GenBank (accession numbers PZ464491-464538). The maps of the cryo-EM reconstruction are available at the Electron Microscopy Data (EMD) Bank under accession number EMD-77188, EMD-77189 and EMD-77190 for H9 HA with batFab30, batFab31 or batFab43 respectively. Coordinates for the complexes are available at the Protein Data Bank under accession number 35UD, 35UE and 35UF for H9 HA with batFab30, batFab31 or batFab43 respectively. The raw movies are available on the electron microscopy public image archive (EMPIAR) under accession number EMPIAR-13579, EMPIAR-13580 and EMPIAR-13581 for H9-batFab31, batFab30 and batFab43 respectively. All other data needed to support the conclusions of the paper are present in the paper or the Supplementary Materials. Requests for material should be addressed to Aaron G. Schmidt.

## MATERIALS AND METHODS

### Approvals, ethics, and biosafety

Bat trapping and sample collections were conducted under permits from the Kenyan National Commission for Science, Technology and Innovation (permit no. NACOSTI/P/18/76501/22243) and the Kenya Wildlife Service (permit no. KWS/BRM/500), University of Nairobi Faculty of Veterinary Medicine; Biosafety, Animal use and Ethics committee (REF: FVM BAUEC/2018/180). Sample import to Finland was approved by the Finnish Food Safety Authority (2809/0460/2018).

### Sample collection

Bats were captured in Taita-Taveta county, Kenya, using mist nets positioned near water sources. They were carefully removed from nets by trained scientists, placed into standard individual cotton bat bags, and transported to an outdoor processing area at the nearby University of Helsinki Taita Research Station. There they were euthanized using an overdose of inhalation isoflurane, followed by cervical dislocation, and dissected for the collection of organ samples. Leather gloves were used to handle live bats, and appropriate PPE was used to process bats, including double layer latex gloves, safety gowns, and FFP3 face masks.

### RNA sample collection and preparation

Spleen samples were placed into separate marked tubes with RNAlater (Sigma), stored at −20°C, and later sent on dry ice to Helsinki, Finland. At the University of Helsinki, under enhanced BSL-3 conditions, bat tissue samples were treated with Tripure (Roche) to inactivate any potential hazardous agents before RNA extractions according to the Tripure protocol. The extracted RNA was stored in −80°C until shipment on dry ice.

### 5’-RACE

5’-RACE protocol was performed using the SMARTer RACE 5’/3’ kit (Takara Bio). Total splenic RNA from each bat was digested with ezDNAse (Thermo Fisher, ezDNase Enzyme) and immediately used as starting material for first-strand cDNA synthesis. First-strand cDNA synthesis was performed in accordance with manufacturer’s protocol using 10X random primer mix (1 µM), 5’ CDS Primer A (0.6 µM), and SMARTer II A oligonucleotide (1.2 µM) in a 20 uL reaction volume, which contained 1 µg of DNAse-treated total RNA, 1x first-strand buffer, RNAse inhibitor (1U) and reverse transcriptase (10U). Primers were annealed at 72°C for 3 min, 42C for 2 min and PCR extension was performed at 42°C for 90 min and inactivated at 70°C for 10 min. Resulting 5’ RACE-ready cDNA was diluted in 90 µL tricine-EDTA buffer and stored at −20°C.

Immunoglobulin heavy chains (IgG and IgM) and light chains (kappa and lambda) were selectively amplified from total first-strand cDNA with universal forward primer (o uniRACE_fw, 0.2 µM) and chain-specific reverse primers (oIgG_RACE_rev, oIgM_RACE_rev, okappa_RACE_rev, olambda_RACE_rev, 0.2 µM) in a 50 µL reaction volume, which contained 2.5 uL 5’-RACE ready cDNA, 1x SeqAmp buffer, and SeqAmp DNA polymerase (Takara Bio). Primers were designed with partial Illumina adapters for downstream sequencing. Thermocycler conditions were optimized to (94°C 30 s, 72°C (−1°C/cycle) 30 s, 72°C 2 min) x 5 cycles, (94°C 30 s, 68°C 30 s, 72°C 2 min) x 25 cycles, 12°C hold. 10 ng of each amplicon (IgG, IgM, kappa, lambda) was diluted to 20 µL volume and indexed with 5 µL of NEBNext unique dual index primers (ERB2: P64-F4, ERB3: P65-F5), 13 µL water, 1 µL dNTP, 1 µL Q5 polymerase, and 10 µL Q5 buffer. The cycling conditions were 98°C for 30 s; 12 cycles of 98°C for 10 s, 72°C for 60 s; and 72°C for 5 min. PCR amplicons were pooled and run on a 1% agarose gel. Amplicons were extracted and purified, then re-purified using AMPure XP beads (Beckman). Amplicons were analyzed by a D1000 TapeStation, quantified with Qubit, and submitted for sequencing on an Illumina MiSeq (2×300 bp kit, Bauer Core).

### Degenerate framework 1 and 4 primer design and analysis

Framework 1 and 4 degenerate primer sets were designed using ERB BCR sequencing from this study. These primer sets were designed to amplify the variable regions from heavy chain and light chain immunoglobulin transcripts without using existing Ig discovery tools that could bias discovery for genes based on sequence similarity to characterized Ig genes. Illumina MiSeq (from above) paired reads were merged with PEAR^81^, demultiplexed by bat, and filtered for reads greater than 500bp that contain the universal forward primer sequence. Reads were then categorized as IgM, IgG, kappa, or lambda based on the presence of the respective reverse primer sequences. Next, for each amplicon, all three reading frames were translated and used to generate a list of all possible ORFs longer than 50 amino acids. Each reading frame was then analyzed with SignalP-6.0 in fast/eukarya mode. For each amplicon, the ORF with the greatest signal peptide and cut site probabilities was selected as the most likely correct ORF, with a minimum accepted probability of 90%. Next, for each of heavy chains, kappa chains, and lambda chains, the first 10 amino acids after the cleavage site were hierarchically clustered with centroid linkage and flat clusters extracted with a distance of 3. We found that visually, the clusters were similar sequences, but misaligned based on small deviations in the annotated cut sites. We assembled a custom database (separately for heavy, kappa, and lambda chains) of all unique post-signal sequence 10 amino acid sequences from clusters with >100 members and manually aligned the clusters together to identify the correct cut site. The original ORF sequence for each amplicon was then searched against its respective heavy, kappa, or lambda database, allowing up to one amino acid mismatch, to annotate the N-terminus of the variable region. The C-terminus was identified via the conserved constant regions: ATTKAPS for IgG, ESSSAPV for IgM, RDVAKP for IgK, and GQPKAGPS for IgL, allowing for a single amino acid mismatch for IgG, IgM, and IgK and two mismatches for IgL. These sequences represented the filtered variable gene list for meta-analysis and downstream primer design.

For primer design, our goal was to design as few degenerate primers as possible for each of IgG/M-forward, IgG/M-reverse, IgK-forward, IgK-reverse, IgL-forward, and IgL-reverse to capture >75% of sequences. The first and last 20 nucleotides of each variable region was isolated, hierarchically clustered with centroid linkage, flattened with a distance metric *t*, and degenerate primers designed for clusters with over *z* members. Metrics *t* and *z* were tuned manually for each primer to keep the number of primers as low as possible while capturing as many variable gene sequences as possible. We found that in some cases clusters were similar enough that the resulting primers could be merged together by introducing one or two additional degenerate nucleotides. Within a cluster, at each position, a nucleotide was included if it was present in >10% of sequences in that cluster at that position. IgG/M-forward: *t*=3.4, *z*=100 resulted in six clusters; IgG/M-reverse: *t*=3.1, *z*=100 resulted in four clusters; IgK-forward: *t*=4, *z*=1000 resulted in two clusters; IgK-reverse: *t*=4, *z*=100 resulted in five clusters; IgL-forward: *t*=4, *z*=1000 resulted in seven clusters; IgL-reverse: *t*=4, *z*=100 resulted in four clusters. Final primer sequences are listed in Supplementary Table 4.

### Bat scFv insert generation

Heavy chain and light chain primer mastermixes were created by mixing equimolar amounts of each designed degenerate primer at 10mM final concentration. The variable region was amplified separately for each bat, keeping IgG, IgM, kappa, and lambda in separate reactions.

Heavy chain and light chain degenerate primers were mixed at equimolar amounts to a final concentration of 10 uM following the compositions in the table above. The heavy chain variable regions (V_H_) were amplified separately from IgG and IgM 5’RACE products using 5µL of oVH_fw and 5µL of oVH_rev for a final primer concentration of 0.5µM. The light chain variable regions (V_L_) were amplified from kappa and lambda 5’ RACE products using 5µL of either kappa or lambda primer mastermix at a final concentration of 0.5 µM. 2 ng of each 5’ RACE product was used as template and reactions were set up using Q5 high-fidelity DNA polymerase (NEB) using cycling conditions 94°C, 5 min, (94°C 30 s, 50°C 30 s, 72°C 30 s) x 20 cycles, 72°C 3 min, 12°C hold. PCR products were separated on a 1% agarose gel, the ∼350bp band was gel extracted and purified (Buffer QG, Qiagen and NucleoSpin Gel and PCR clean-up columns, Macherey-Nagel).

Overlap PCR was used to link V_H_ and V_L_ products with a 6x Gly-Ser linker to create scFv inserts. Inserts have homology overhangs with yeast display plasmid pCHA^82^. 40 ng of VH and VL each was used to generate V_H_-kappa and V_H_-lambda scFvs, with 0.5 µM of oOE_fw and 0.5 µM of oOE_rev using Q5 high fidelity polymerase (NEB). Cycling conditions were 94°C 5 min, (94°C 30 s, 70°C 30 s, 72°C 1 min) x 10 cycles, 72°C 4 min, 12°C hold. PCR products were separated on a 1% agarose gel, the ∼750bp band was gel extracted and purified (Buffer QG, Qiagen and NucleoSpin Gel and PCR clean-up columns, Macherey-Nagel). Overlap extension PCRs were amplified using oOE_fw and oOE_rev to generate insert for yeast electroporation.

### Yeast display library generation

Yeast cells (strain AWY101) were electroporated with 5µg of scFv insert and 1µg of digested pCHA plasmid as previously described^49^. To maximize library size, 5 libraries of V_H_-kappa scFv (estimated diversity ∼7.2E7 per electroporation, ∼3.6E8 combined kappa diversity) and 5 libraries of V_H_-lambda scFv (realized diversity of ∼6.2E7 per electroporation, 3.1E8 combined lambda diversity)) were electroporated, which were pooled for a theoretical library size of ∼6.7E8 with approximately equal representation of lambda and kappa scFvs from each bat.

### Identification of putative V genes

823981 paired heavy chain reads were mapped to partial (estimated ∼90% coverage) ERB genome Raegyp2.0^31^ using BLAST 2.17.0. loci. Paired reads mapped were filtered to include only those that were at least 250 bp in length and that contained a known ERB recombination signal sequence within 30 bp downstream of the end of the mapped region^12^. All sequences were visually inspected to exclude those that appeared to be pseudogenes or contained internal truncations, deletions, or stop codons. To identify the precise start and end position of each putative V_H_ gene, mapped sequences were extracted from Raegyp2.0 including an additional 30 bp both upstream and downstream; V_H_ genes boundaries were identified from these sequences using IMGT/LIGMotif^83^.

### Genetic analysis of ERB antibody transcripts

A database of germline V, D, and J genes was assembled using all functional genes previously described^12^ in addition to all putative V_H_ genes described above. IgBLAST 1.17.1^84^ was used with the resulting database to map and determine the percent identity of each IgM and IgG sequencing read to its nearest germline ERB V/D/J gene, along with identifying the CDR3 region. For kappa and lambda sequencing reads, for which there are no germline ERB databases, human kappa V/J and lambda V/J genes obtained from IMGT^85^ as reference.

### Somatic hypermutation (SHM) frequency

To calculate frequency of mutation from germline at each position in the IGHV gene, IgBLAST 1.17.1 was used to map each IgM and IgG sequencing read to its nearest germline IGHV gene. To avoid overestimating mutation frequency as a consequence of some reads being incorrectly mapped to a germline gene, only those sequences with at least 80% germline identity were retained. MUSCLE5’s Super5 multiple sequence alignment (MSA) algorithm^86^ was then used to produce a single MSA of all germline IGHV genes and paired reads together. All MSA nucleotide positions determined to be a gap were ignored for all germline sequences. At every remaining MSA position, mutation frequency was defined at that position to be the proportion of sequencing reads not having the same nucleotide (or gap) as their parent germline gene at said position.

### Expression and purification of recombinant influenza antigens

Plasmids encoding the FLsE of group 1 and group 2 influenza HA were designed for H1N1 A/Solomon Islands/3/2006, H2N2 A/Japan/305-/1957, H3N2 A/Hong Kong/1/1968, H5N1 A/Vietnam/1194/2004, H7N9 A/Shanghai/02/2013, H9N2 A/Bat/Egypt/381OP/2017, H9N2 A/Guangdong/MZ058/2016, H17N10 A/little yellow-shouldered bat/Guatemala/060/2010, H18N11 A/flat-faced bat/Peru/033/2010. Native signal peptide (predicted by SignalP-6.0) and transmembrane domain (predicted by TMHMM-2.0) were removed for full length HA secretion. Head-only construct for H9 A/Bat/Egypt/381OP/2017 was designed with residues 42 to 324; H9 numbering. Constructs were codon optimized and synthesized by Integrated DNA Technologies (IDT) and cloned into pVRC8400 protein expression vectors with C-terminal tags: H1, H2, H3, H7 FLsE Foldon (Fd) trimerization tag, 8xHis, and streptavidin binding protein (SBP) tags); H5, H9 GD16, H9 head-only (Fd, 8xHis, and Avi tags); H17 FLsE (GCN4 trimerization tag, HRV 3C-cleavable 8xHis, Avi tags); H9 EG17 and H18 FLsE with universal influenza stem stabilizing mutations and fusion loop R to A mutation (HRV 3C-cleavable 8xHis, Avi tags)^51^. Proteins were transiently expressed in Expi293F cells (Thermo Fisher Scientific). Five to seven days after transfection, supernatants were harvested by centrifugation and purified using cobalt-TALON resin (Takara) followed by size exclusion chromatography on Superdex 200 Increase 10/300 GL column (GE Healthcare).

### Probe generation

Avi-tagged FLsE were biotinylated using BirA biotin-protein ligase kit (Avidity) according to the manufacturer’s protocol and subsequently re-purified by size exclusion chromatography on a Superdex 200 Increase 10/300 GL column (GE Healthcare).

### Enrichment for antigen-specific yeast clones

Yeast sorting was performed as previously described using at least 10-fold more yeast than the number of cells isolated from the previous round of enrichment.^49^. Briefly, the bat scFv library was grown in SDCAA media at 30°C with shaking (250 rpm) and surface expression of bat scFvs was induced in SG-CAA media for 36 hours at 20°C with shaking (250 rpm). Candidate H9-reactive clones were enriched over two rounds of magnetic activated cell sorting using a final concentration of 1 μM biotinylated H9 HA with streptavidin microbeads (round 1) and anti-biotin microbeads (round 2) (Miltenyi). For fluorescence activated cell sorting (FACS) based enrichment, cells were isolated using two-color labeling for expression (chicken anti-c-Myc IgY (Invitrogen, 1:250); Alexa Fluor 488-donkey anti-chicken IgG (Invitrogen, 1:200) and binding (streptavidin-allophycocyanin (Invitrogen, 1:100), neutravidin-phycoerythrin (Invitrogen, 1:50), or anti-biotin IgG-allophycocyanin (Miltenyi, 1:50)), as previously described^49^. In each round of FACS, biotinylated H9 HA was titrated to an appropriate concentration to enrich for tight binders and staining volume was adjusted so that moles antigen remains in excess of available scFvs. In R2 FACS, biotinylated H9 HA was titrated from 1μM to 250 nM and 100 nM and binders were sorted. Forty-eight single yeast clones were sanger sequenced, twenty-four of which were also characterized on FACS for binding against 100 nM H9 HA. The DNA plasmid was extracted from polyclonal yeast cells reactive against 100 nM H9 HA and scFvs were PCR amplified and deep sequenced on the minION Mk1B using a minION flow cell (Oxford Nanopore Technologies). A secondary-only control was used to isolate yeast clones nonspecifically sticking to the secondary, leading to a false positive. DNA plasmid was similarly extracted from the non-specific polyclonal yeast cells, PCR amplified, and deep sequenced on the minION Mk1B using a Flongle flow cell (Oxford Nanopore Technologies). Nonspecific sequences were removed from sequencing analysis. H9 HA was titrated to 1 nM in R3 and R4 of FACS, toggling secondaries to prevent enrichment of nonspecific fluorophore binders. The plasmid DNA was extracted from polyclonal yeast cells after R4 of FACS, along with nonspecific yeast clones as described above. The scFvs were PCR amplified from the plasmid DNA, barcoded, and deep sequenced on the PromethION 2 using a PromethION flow cell (Oxford Nanopore Technologies). Nonspecific sequences were removed from sequencing analysis.

### Genetic analysis of H9 HA-reactive yeast clones

To address the high sequencing error rate of Oxford Nanopore sequencing, only sequences that appeared more than ten times were considered in analyses as it is highly unlikely for the same random sequencing error appearing more than ten times. Nonspecific sequences were removed from the 100 nM H9 HA-reactive and 1 nM H9 HA-reactive datasets so that analyses were performed on H9 HA-reactive sequences only. The genetic analyses performed on these antibody sequences were identical to those described above that were applied to the Illumina ERB antibody repertoire sequence datasets.

### Expression and purification of monoclonal bat antibodies

Bat IgG genes for heavy- and light-chain constant domains were synthesized, codon optimized and subcloned into pVRC8400 protein expression vectors based on published bat constant domain sequences^12^. Bat IgG1 plasmids contain a C-terminal HRV 3C cleavable 8xHis tag. The sequenced heavy and light chain variable regions from H9 reactive yeast clones were synthesized by IDT and subcloned into the IgG and light chain constant plasmids described above using Gibson Assembly (NEB). IgGs were similarly expressed and purified above for influenza HAs, and buffer exchanged into phosphate-buffered saline (PBS). Amino acid sequences of the recombinantly expressed batAbs are provided in data file S2.

### Enzyme-linked immunosorbent assay

BatAb reactivity to HA antigens were assayed by ELISA. Briefly, 96-well high-binding (Corning) microplates were coated with HA (4 μg/mL) in PBS at 100 μL per well and incubated overnight at 4°C. Plates were blocked with 1% bovine serum albumin (BSA) in PBS containing 1% Tween 20 (PBS-T) for 1 hour at room temperature (RT). Blocking solution was removed and sixfold serial dilutions of bat IgG1 (50 μg/mL starting concentration) in PBS were added to wells and incubated for 1 hour at either RT. Plates were washed three times with PBS-T. Protein A/G-horseradish peroxidase (HRP; Pierce) was added at 1:10,000 dilution in PBS for 1 hour at RT. Plates were washed three times with PBS-T and developed with 1-step ABTS substrate (Pierce).

Absorbance was measured using a plate reader at 450 nm. To determine the EC_50_ values of binding recombinant batAbs, ELISAs were repeated as above in triplicate with tenfold serial dilutions of batAbs (300 μg/mL starting concentration) in PBS. EC_50_ values were determined by nonliner regression (sigmoidal) using GraphPad Prism 10.1.0 software. ELISAs to establish HA binding breath were done at a single IgG concentration (50 μg/mL) in triplicate. Positive binding was defined by an optical density at 450 nm (OD_450_) ≥ 0.50.

Streptavidin ELISA plates were coated with recombinant streptavidin (Invitrogen) at 4 μg/mL in PBS as described above. ELISAs were performed in duplicate with tenfold serial dilutions of batAbs (300 μg/mL starting concentration) in PBS.

Polyreactivity ELISA plates against human insulin (MilliporeSigma), dsDNA (calf thymus DNA; Invitrogen), and BSA (Sigma-Aldrich) were coated with 2, 50, and 5 μg/mL, respectively, in PBS and incubated overnight at 4°C. Plates were blocked and incubated with IgGs as described above. Cardiolipin ELISAs were measured as previously described^87^. Briefly, plates were coated with 5 μg of cardiolipin per well (from bovine heart; MilliporeSigma, C1649) in 30 μL ethanol uncovered overnight at 4°C. Plates were washed twice with 100 μL per well of 1x PBS and blocked with 100 μL PBS with 10% FBS for 1 hour at RT (covered). Blocked plates were washed three times with 1x PBS and incubated IgGs were diluted in 1x PBS with 1% FBS for 1 hour at RT. After washing with 1x PBS, cardiolipin ELISAs were developed as described above. Polyreactive ELISAs were performed at a single IgG concentration (50 μg/mL) in triplicates, and polyreactivity was defined as (OD_450_) ≥ 0.40.

### Cryo-EM sample and grid preparation

Cryo-EM grids for H9-Fab43 complex were prepared as follow, H9 (2.0 mg/mL) was mixed with Fab43 at a 1:1.2 molar ratio and incubated on ice for 1 hour, immediately before applying the sample to the grid, n-Octyl-β-D-Glucoside (Anatrace, O311) was added to the preformed complex at 0.1% w/v final concentration. Sample was applied on glow discharged (15 mA, 25 s) Quantifoil® R0.6/1 grids (Ted Pella Inc, N1-A11nAu30-50) and plunge-frozen into liquid ethane using a Vitrobot Mark IV (Thermo Fisher Scientific) with a blotforce of 0 and a blot time of 4-5.5s under chamber conditions maintained at 4°C and 100% humidity. H9-Fab30 and H9-Fab31 complexes were prepared under identical conditions and samples were applied to glow-discharged Ultrathin Carbon on Quantifoil® 1.2/1.3 grid (Ted Pella Inc, 668-400-AU). Initial screening of grids was performed on a 200 kV Glacios (Thermo Fisher Scientific) at Icahn school of Medicine at Mount Sinai cryo-EM core facility. Grids exhibiting optimal ice thickness and particle distribution and orientation were saved for high-resolution data collection either at MSSM cryo-EM core or NYSBC.

### Cryo-EM data collection

The H9-Fab30 dataset collected at NYSBC comprised 8,758 movies collected on a 300 kV Titan™ Krios (Thermo Fisher Scientific) equipped with Gatan K3 direct detector, and BioQuantum imaging filter. Data w^60^ere acquired at a physical pixel size of 0.856Å, with a total dose of 55.67 e−/Å2. The H9-Fab31 along with the H9-Fab43 dataset, were collected at MSSM cryo-EM Core on a Glacios™ G2 (Thermo Fisher Scientific) equipped with a X-FEG electron gun and Falcon IVi detector. A total of respectively 6,395 and 10,430 movies were acquired at a pixel size of 0.722 Å, with a total dose of 50 e−/Å2. Full data collection parameters are summarized in **table S3**.

### Cryo-EM data processing

#### H9 HA-batFab30 Complex

Movies were aligned and dose weighted using patch motion correction implemented in cryoSPARC V4.7.1^60^. Contrast transfer function estimation was done in cryoSPARC V.4.7.1 using Patch CTF. After blob picking and a first round of 2d classification, 2D classes corresponding to multiple projection of the particles were selected and used as template for template picking. The picked particles were extracted with a box size of 512 pixels, with 4x binning, and subjected to a 2D classification. After discarding particle images corresponding to obvious junk, the rest of the particle images (1,149,347) were used to generate 3 initial model. The particles originating the best volume (54%) underwent 3D non-uniform refinement followed by one round of unsupervised 3D classification without alignment. Particle stack associated with the best occupancies and overall map quality were selected for one round of heterogeneous refinement using as input the map with 2 fab, the map with 3 fab and a decoy map corresponding to previously discarded particles from the initial ab initio job.

Particle images corresponding to volume representing HA harboring 2 fabs was further processed without symmetry imposed. After one round of 3D classification, heterogeneous refinement job cloned from previously described, local and global CTF refinement and per-particle motion correction and final round of 3D NUR, 91,645 particles yielded a final map with nominal resolution of 3.0Å based on the gold-standard Fourier shell correlation od 0.143 criterion. Particles corresponding to volume of HA interacting with 3 Fabs were subjected to one round of Ab initio followed by 3D NUR, local and global CTF refinement, per-particle motion correction and final round of 3D NUR with C3 symmetry imposed, 209,939 particles yielded a map at a nominal resolution of 2.35Å based on the gold-standard Fourier shell correlation od 0.143 criterion.

#### H9 HA-batFab31 Complex

Movies were aligned and dose weighted using patch motion correction implemented in cryoSPARC V4.7.1. Contrast transfer function estimation was done in cryoSPARC V4.7.1 using Patch CTF. After blob picking and a first round of 2D classification, 2D classes corresponding to multiple projection of the particles were selected and used as template for template picking. The picked particles were extracted with a box size of 512 pixels, with 4x binning, and subjected to a 2D classification. After discarding particle images corresponding to obvious junk, the rest of the particle images (599,538) were used to generate 5 initial model. The particles originating the best volume (44%) underwent 3D heterogeneous refinement (best map + 2 bad maps), best particles underwent ab initio, 3D heterogeneous refinement and finally 3D NUR (strategy based on CS automated workflow). Particles were sieved using cryoSIEVE^88^ and a stack of 59,533 was finally re-imported to Cryosparc, particles were refined 3D NUR followed by local and global CTF refinement, per-particle motion correction. Followed by a final round of 3D NUR with C3 symmetry imposed, yielding a map at a nominal resolution of 2.90Å based on the gold-standard Fourier shell correlation od 0.143 criterion.

#### H9 HA-batFab43 Complex

Movies were aligned and dose weighted using patch motion correction implemented in cryoSPARC V4.7.1. Contrast transfer function estimation was done in cryoSPARC V4.7.1 using Patch CTF. After blob picking and a first round of 2D classification, ∼1M particles were used to generate 2 initial volumes. After 2 rounds of heterogeneous refinement with 3 model volumes (1 good and 2 bad) the best volume was used to generate templates for 2D template picking. After further curation a consensus map corresponding to 807,433 particle was generated (3D NUR, local/global CTF refinement and per-particle motion correction yielding a map at a nominal resolution of 2,74Å.This stack of particle was further classified by 3 rounds of heterogeneous classification using 1 good and 2 bad volumes as templates and only the particles associated to the map with the best Fab occupancy were subjected to the following round of 3D heterogeneous refinement (as input for good volume the volume corresponding to the map with the Fab). The final stack was the imported to cryoDRGN for further heterogeneity analysis. A stack of ∼30,000 particles associated to a map with a high Fab occupancy were reimported to CS and refined to generate a template and used to train topaz for picking along with the initial stack imported to cryoDRGN. Particles picked by Topaz were subjected to rounds of 2D classification (strategy based on 2D template matching^89^) followed by multiple rounds of ab initio and heterogeneous refinement^90^. Particles stack associated with best defined Fab along with particles re-imported from cryoDRGN were further classified using unsupervised 3D classification (k=40).

### Model Building and Refinement

The DeepEMhancer sharpened focused maps were used for model building. Model for H9 was initially built using consensus map from H9 HA-batFab43 complex by rigid body fitting of AlphaFold^91^ model prediction followed by multiple rounds of real-space refinement in Phenix^92^ and manually adjustment in Coot^93^. Individual Fab models were generated using alphafold3 server and rigid body fitted in the corresponding density, then the H9 HA-batFab models were subjected to multiple rounds of real-space refinement and manual adjusting using COOT, finally models were validated with MolProbity^94^ and privateer. Figures and structural analysis were done using ChimeraX^95^. The structural biology software was compiled and distributed by SBGrid.

