## Supplemental Information for "Egyptian rousette bat humoral immunity to H9 influenza hemagglutinin"

**A**

| | Total | Unique |
|---|---|---|
| Heavy chain | 823,981 | 823,959 |
| Lambda | 1,285,200 | 1,226,762 |
| Kappa | 1,920,159 | 1,913,188 |

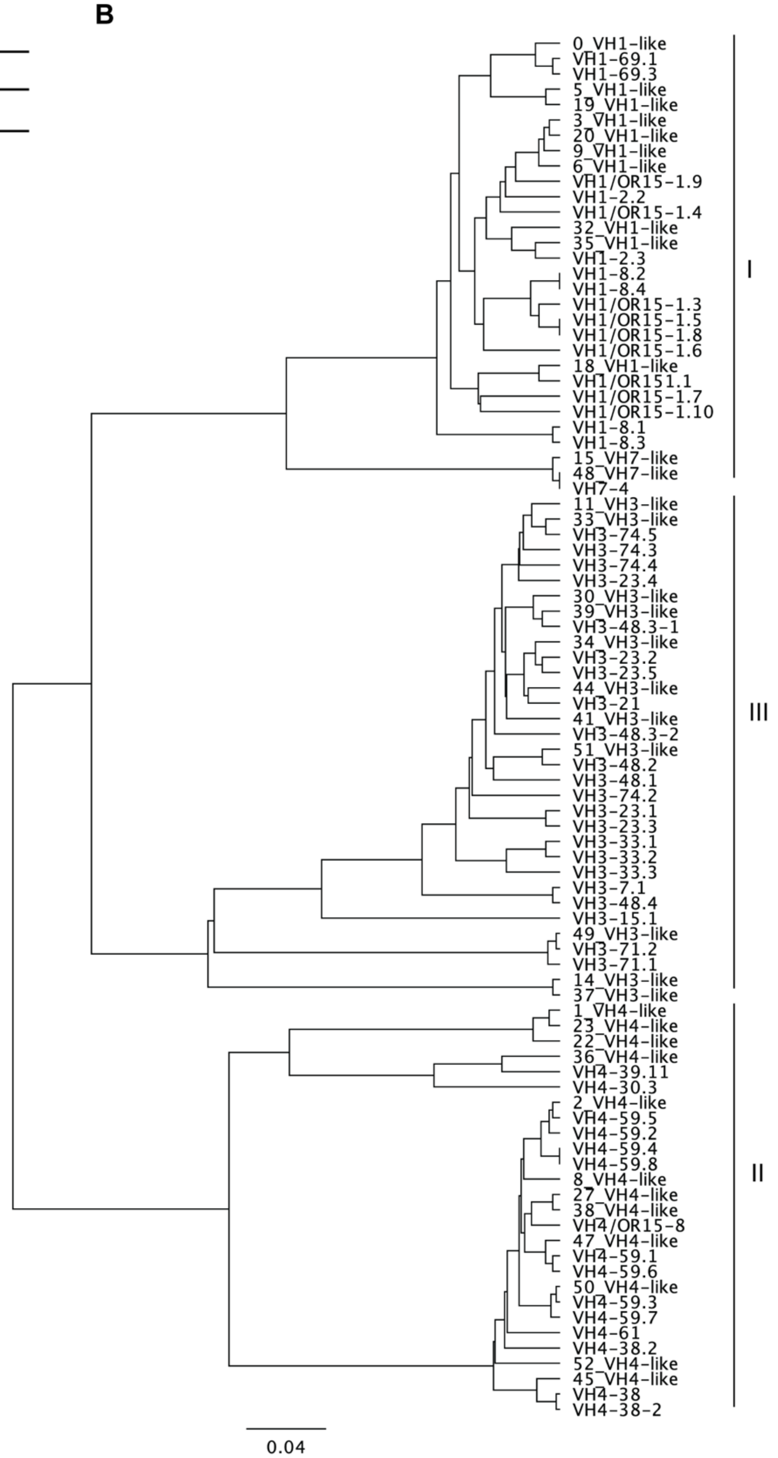


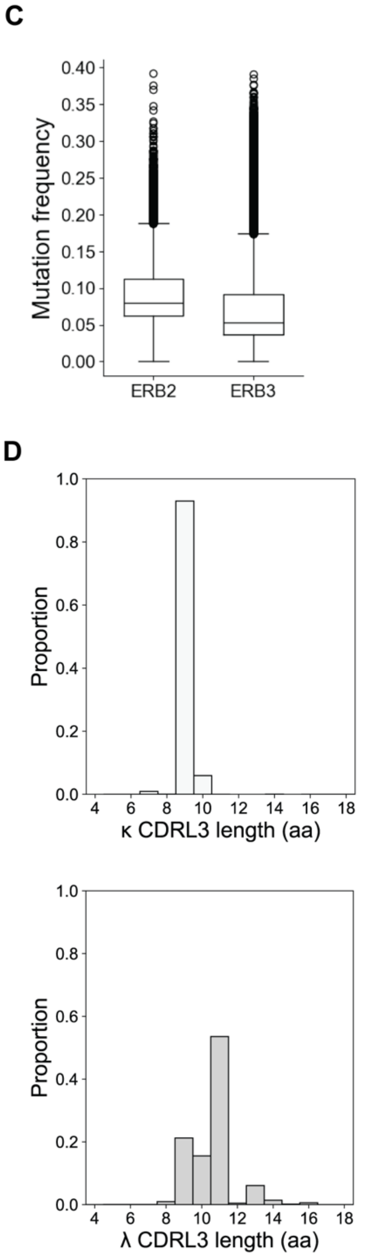


**fig S1. Putative ERB $V_H$ genes. (A)** Counts of total and unique BCR sequences for heavy, lambda, and kappa chains. **(B)** Phylogenetic tree of augmented ERB $V_H$ genes. Tree produced using UPGMA with Geneious Prime 2024.0.7 on $V_H$ nucleotide sequence, excluding leader. $V_H$ genes cluster within defined clans I, II, and III. **(B)** Phylogenetic tree of augmented ERB $V_H$ genes. Tree produced using UPGMA with Geneious Prime 2024.0.7 on $V_H$ nucleotide sequence, excluding leader. $V_H$ genes cluster within defined clans I, II, and III. **(C)** Somatic hypermutation (SHM) frequency of ERB naïve and class-switched repertoires for each bat. Median mutation frequency represented by solid bar. **(D)** CDR L3 length distribution for kappa (top) and lambda (bottom) transcripts.

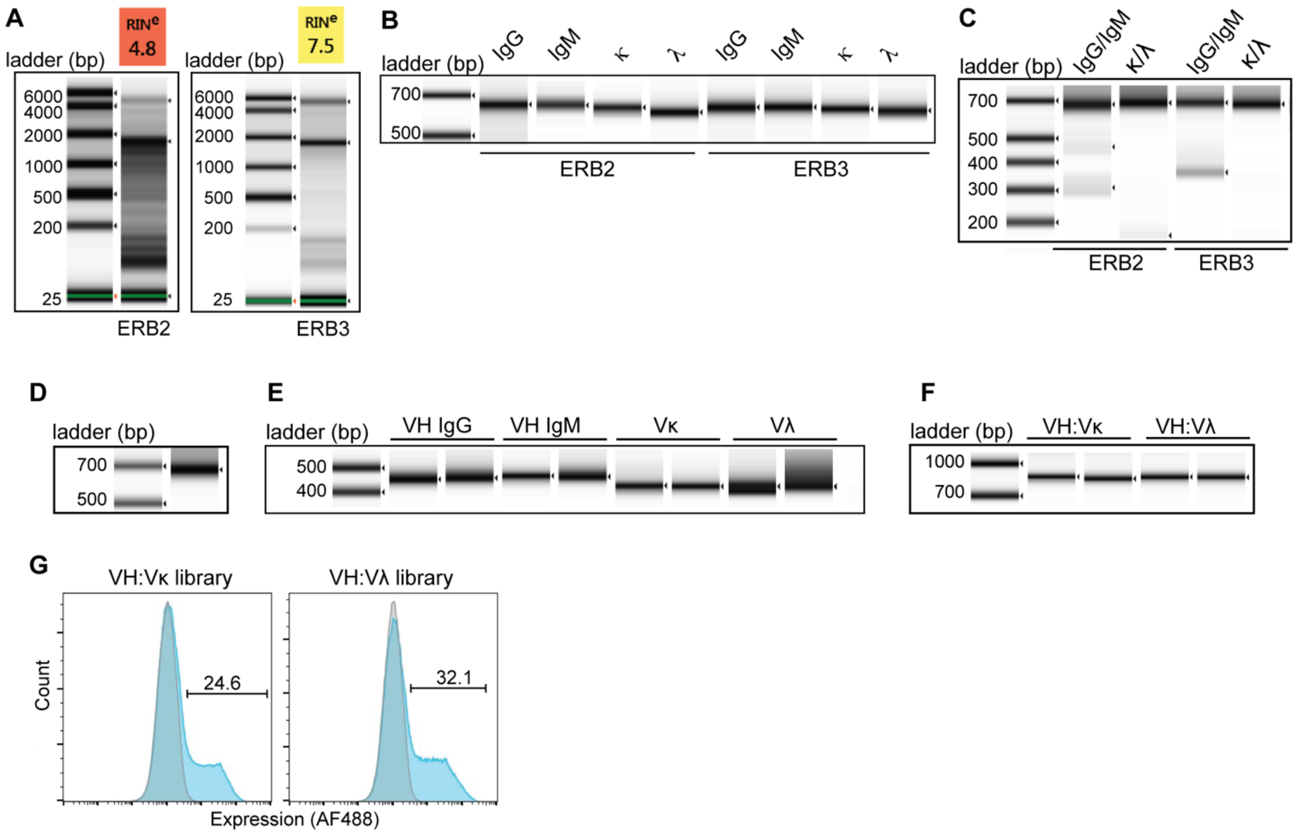


**fig. S2. Library preparation**. **(A)** Total splenic RNA from ERB2 and ERB3. RIN, RNA integrity score. **(B)** DNA D1000 ScreenTape of enriched antibody heavy and light chain transcripts with IgG/M $C_H1$ and lambda/kappa $C_L$ gene-specific primers after bead clean-up. **(C)** NEBNext indexed antibody transcripts prepared for Illumina sequencing. **(D)** Excised and bead cleaned up band of pooled NEBNext indexed transcripts submitted for Illumina sequencing. **(E)** $V_H$ and $V_L$ amplified from 5' RACE cDNA from ERB2 (left) and ERB3 (right) using degenerate framework 1 and framework primer mixes. **(F)** $V_H$-$V_L$ scFv overlap extension PCR for ERB2 (left) and ERB3 (right) prepared for electroporation into AWY101 yeast. **(G)** Expression of $V_H$-Vκ and $V_H$-Vλ on yeast confirmed by flow cytometry detection of C-terminal myc tag.

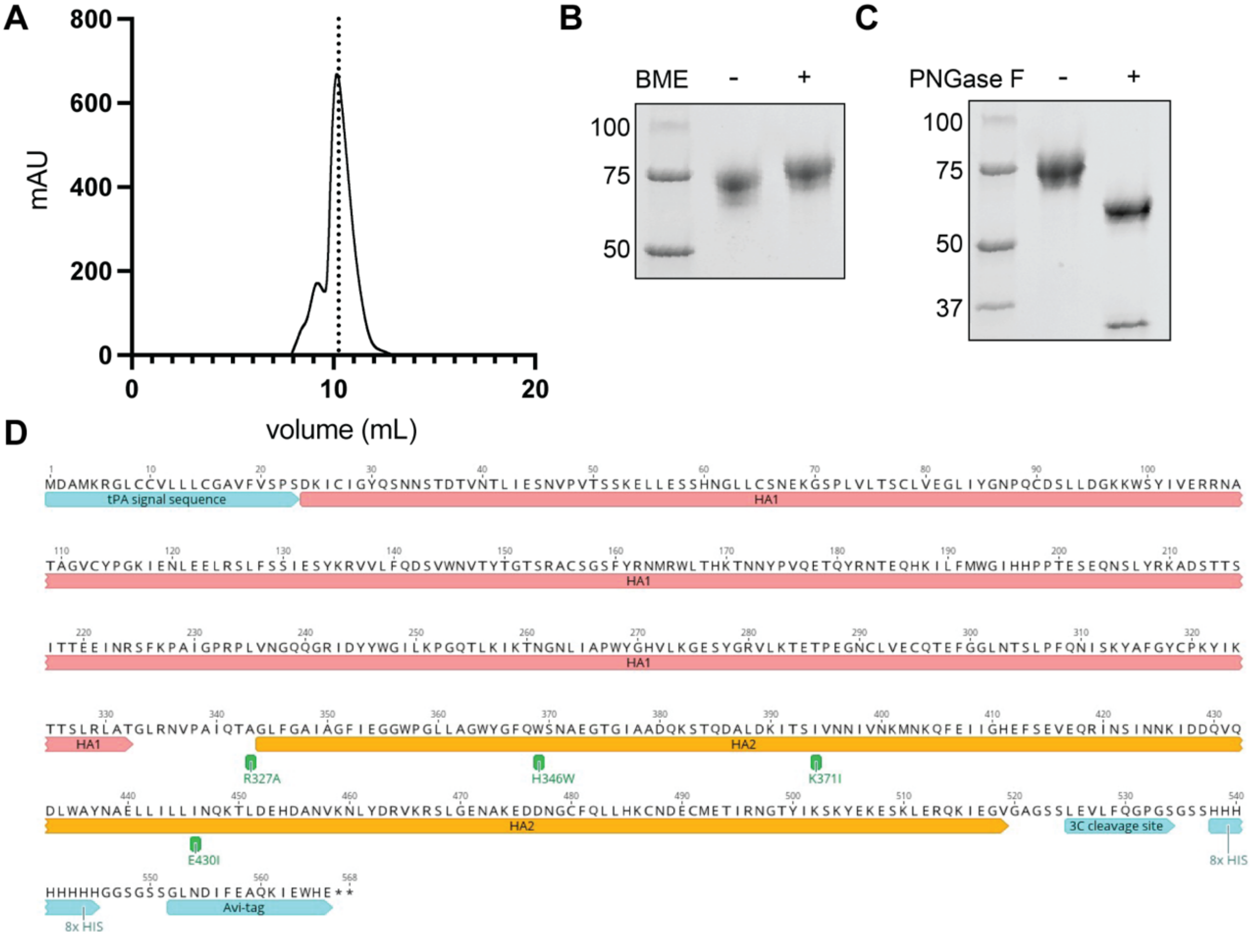


**fig. S3. Bat H9 HA construct design and expression**. **(A)** SEC profile of bat H9 FLsE from a Superdex 200 10/300 column. **(B)** SDS PAGE of H9 FLsE in nonreducing and reducing conditions and **(C)** deglycosylated using PNGaseF (right). **(D)** Annotated H9 HA full length soluble ectodomain (FLsE) with introduced mutations and tags.

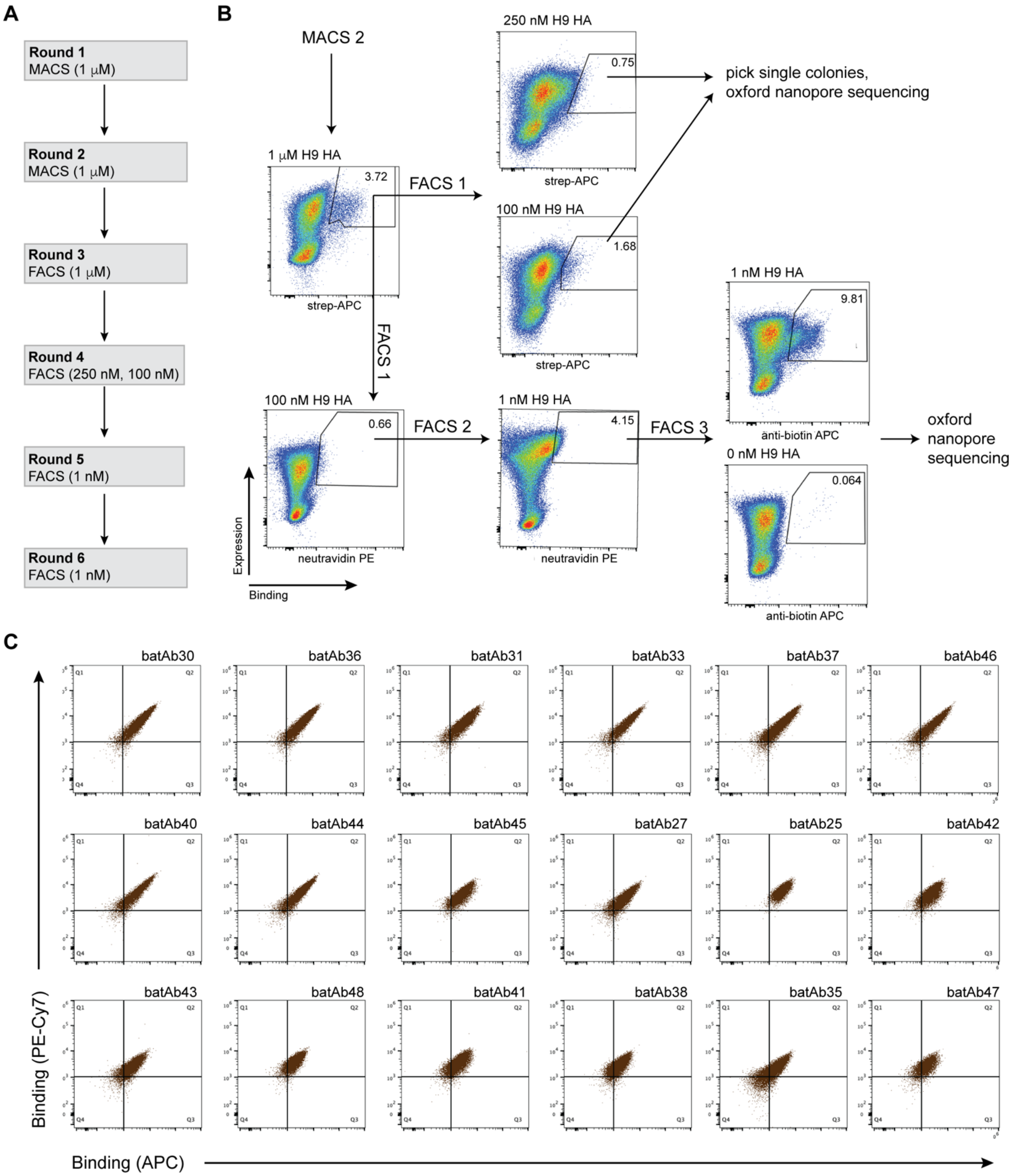


**fig. S4. Enrichment of H9 HA specific batAbs**. **(A)** Selection strategy for batAb enrichment using magnetic activated cell sorting (MACS) and fluorescence activated cell sorting (FACS). H9 HA FLsE:biotin was titrated from 1μM down to 1nM over six rounds of enrichment. **(B)** Flow plots of the full enrichment campaign against H9 HA FLsE:biotin, alternating fluorophores and titrating H9 HA antigen. Gates drawn to indicate sorted populations **(C)** Single clone binding of 100 nM and 250 nM H9 HA reactive batAb yeast clones as indicated by APC and PE-Cy7 signal gated on the expressing population.

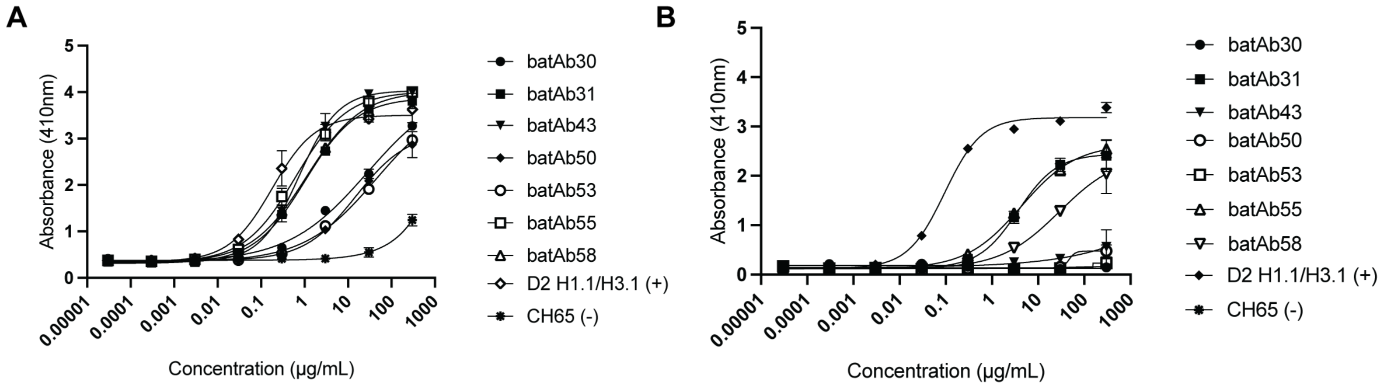


**fig. S5. Recombinant batAb ELISA binding curves.** **(A)** H9 HA full length and **(B)** head-only binding curves performed in technical triplicate. Error bars indicate SD.

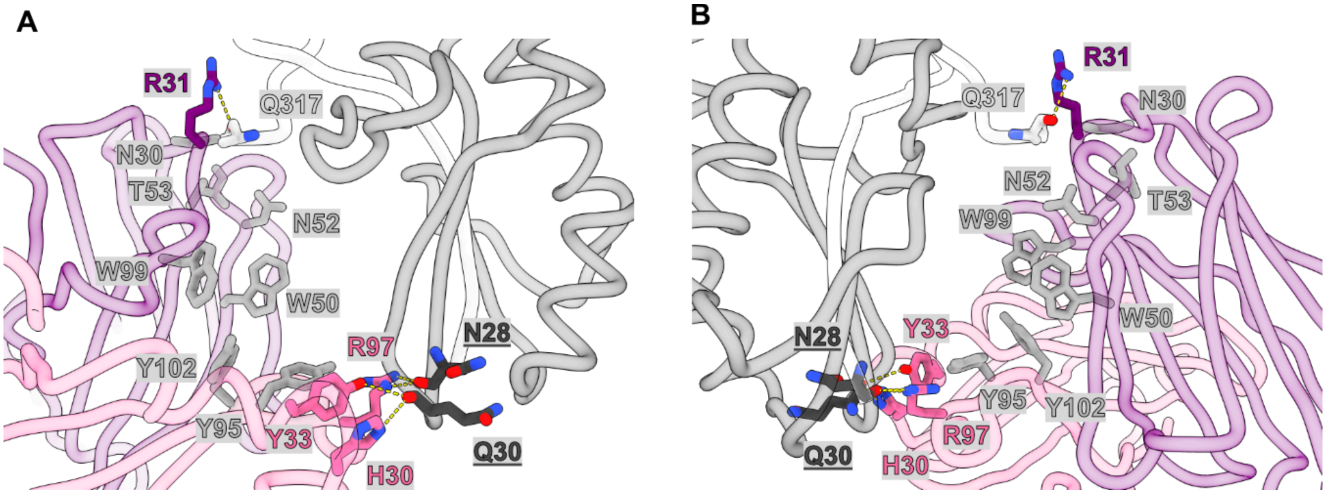


**fig. S6. Superposition of batFab30 with activated H9 HA A/swine/Hong-Kong/9/98.** HA2 domain H9 HA A/swine/Hong-Kong/9/98 (PDB 1JSD) was aligned on the HA2 domain of H9 HA A/Bat/Egypt/381OP/2017 (Residue 351-495, RMSD = 0.798Å) for comparison of footprint of batFab30 between inactivated and activated conformation of H9 hemagglutinin.

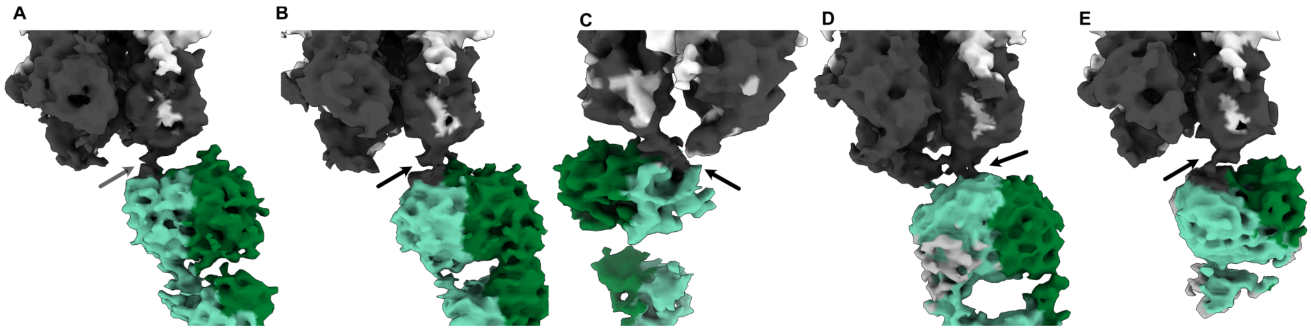


**fig. S7. Cryo-EM maps of the identified discrete orientations of batFab43.** HA1 in white, HA2 in grey, batFab43 light chain in pale green and heavy chain in dark green. The medium resolution maps correspond to maps from **Fig. 4H**. In every map we observe a tube-like volume (dark arrow) from the HA2 C-terminal domain to the light chain of batFab43.

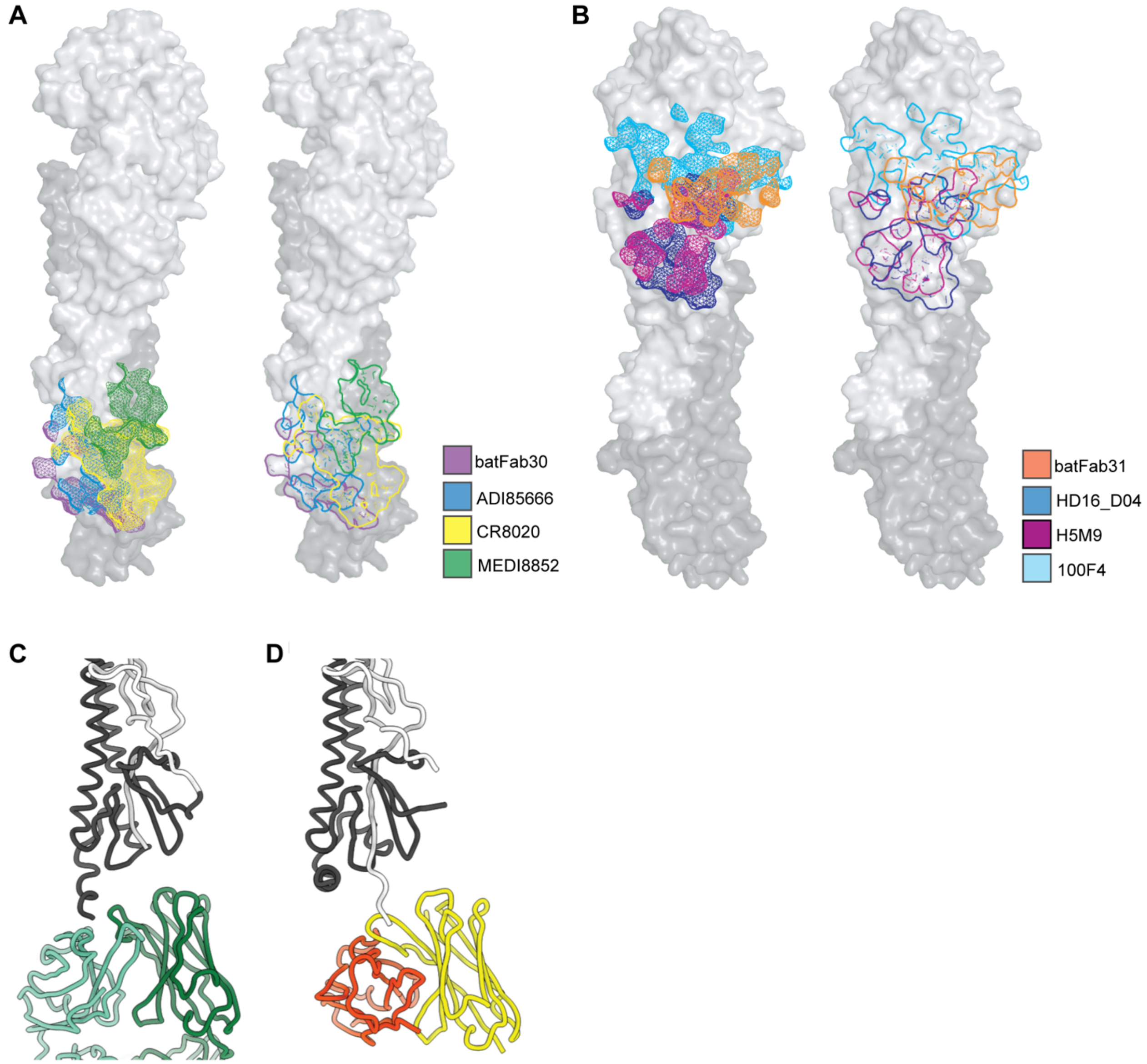


**fig. S8. Footprints comparison of batFab30, batFab31 and batFab43 with known antibodies.** HA1 in dark, HA2 in white. Footprints are outlined on the surface of H9 A/Bat/Egypt/381OP/2017. (**A**) Footprints of batFab30 (purple), ADI85666 (blue), CR8020 (yellow), and MEDI8852 (green) on HA stem fusion peptide domain. **(B)** Footprints of batFab31 (orange), HD16_D04 (blue), H5M9 (purple), 100F4 (cyan) at the vestigial esterase domain. Side to side comparison of batFab43 **(C)** and mouse 5E10 **(D)** binding to H9 HA A/Bat/Egypt/381OP/2017 and H3 HAs A/Victoria/361/2011, respectively. Both HAs were aligned on HA2 (RMSD= 1.183Å).

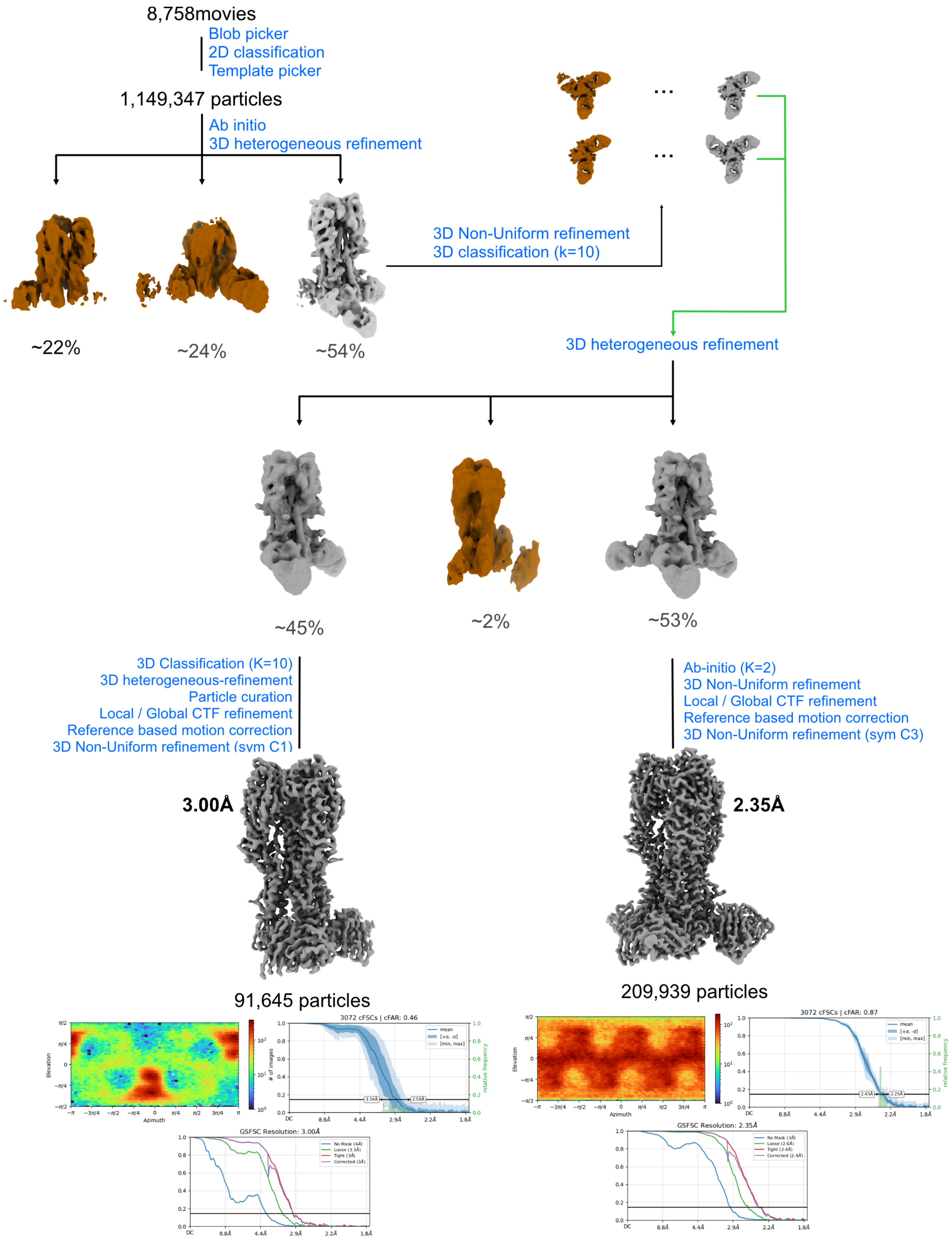


**fig S9. Cryo-EM data processing workflow for determination of the structure of HA A/Bat/Egypt/381OP/2017 in complex with batFab30.**

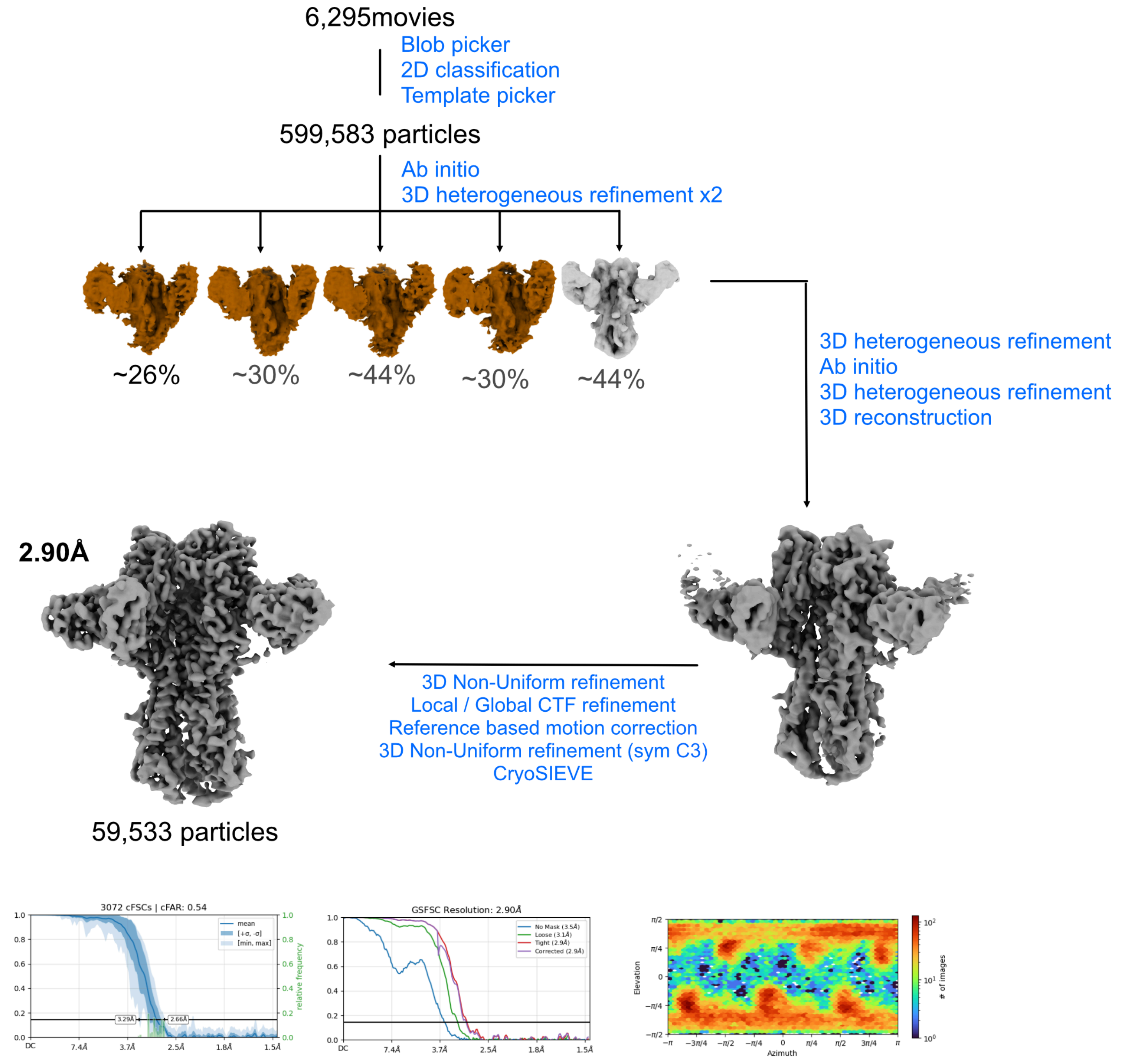


**fig. S10. Cryo-EM data processing workflow for determination of the structure of HA A/Bat/Egypt/381OP/2017 in complex with batFab31.**

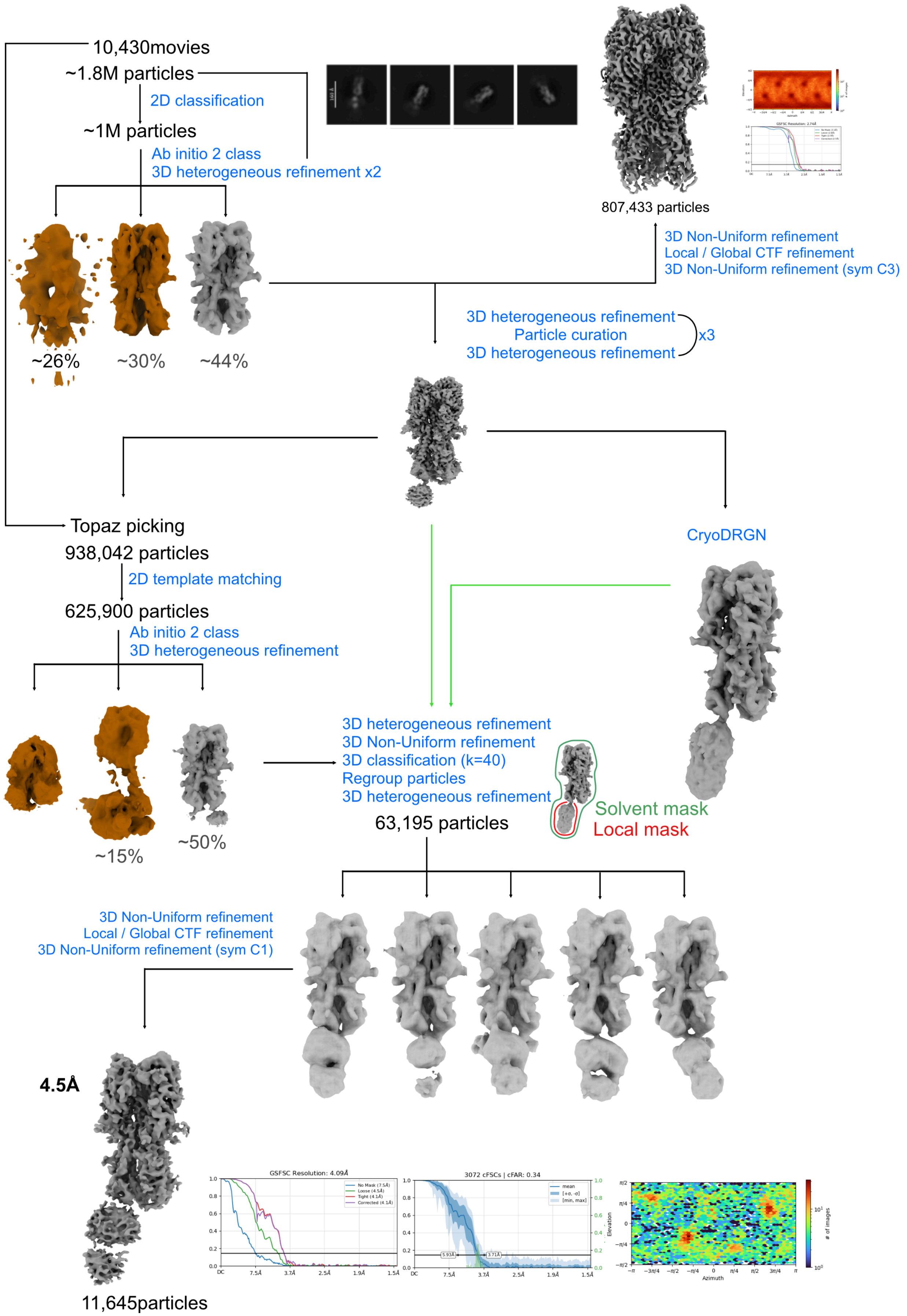


**fig. S11. Cryo-EM data processing workflow for determination of the structure of HA A/Bat/Egypt/381OP/2017 in complex with batFab43.**

**table S1. Putative Egyptian rousette bat $V_H$ genes.** Bolded sequences are putative $V_H$ genes identified in this study.
(see Excel spreadsheet)

**table S2. Recombinant H9 HA reactive bat antibodies.**
(see Excel spreadsheet)

**table S3. Cryo-EM data collection and statistics.**

| Fab:HA A/Bat/Egypt/381OP/2017 complex | | | batFab30:HA | batFab31:HA | batFab43:HA |
|---|---|---|---|---|---|
| Model | Deposition ID | | EMD-77188; PDB 35UD | EMD-77189; PDB 35UE | EMD-77190; PDB 35UF |
| | Composition | Chains | 9 | 12 | 5 |
| | | Atoms (Hydrogens: 0) | 17613 | 16840 | 15037 |
| | | Residues – Protein | 2168 | 2115 | 1922 |
| | | Ligands | BMA: 5, NAG: 31, MAN: 4, BOG: 3 | BMA: 1, NAG: 21, MAN: 1 | N/A |
| | Bonds (RMSD) | Length (Å) (# > 4σ) | 0.005 (0) | 0.005 (0) | 0.004 (0) |
| | | Angles (°) (# > 4σ) | 0.663 (1) | 0.627 (0) | 0.752 (5) |
| | Quality | MolProbity score | 0.73 | 0.97 | 1.16 |
| | | Clash score | 0.63 | 1.02 | 1.24 |
| | Ramachandran plot (%) | Outliers | 0 | 0 | 0 |
| | | Allowed | 2.09 | 2.97 | 4.50 |
| | | Favored | 97.91 | 97.03 | 95.50 |
| | Rama-Z (RMSD) | whole | 0.17 (0.18) | -0.98 (0.18) | -0.64 (0.19) |
| | | helix | 2.30 (0.26) | 1.31 (0.26) | 1.32 (0.27) |
| | | sheet | 0.42 (0.20) | -1.56 (0.22) | -0.24 (0.23) |
| | | loop | -0.82 (0.18) | -0.74 (0.17) | -1.16 (0.18) |
| | Other | Rotamer outliers (%) | 0 | 0 | 0.12 |
| | | Cβ outliers (%) | NA | NA | NA |
| | | Peptide – Cis proline/general (%) | 0.0/0.0 | 0.0/0.0 | 4.9/0.0 |
| | | Peptide – Twisted proline/general (%) | 0.0/0.0 | 0.0/0.0 | 0.0/0.0 |
| | | CaBLAM outliers (%) | 1.03 | 1.89 | 3.21 |
| | ADP (B-factors) | Iso/Aniso (#) | 17613/0 | 16840/0 | 15037/0 |
| | | Protein min/max/mean | 21.28/88.79/44.33 | 32.68/127.07/65.14 | 38.26/1024.55/185.60 |
| | | Ligand min/max/mean | 25.67/105.04/60.71 | 2.36/125.10/83.30 | 0.00/134.27/87.94 |
| | Occupancy | Mean | 1 | 1 | 1 |

| | | | | | |
|---|---|---|---|---|---|
| | | occ = 1 (%) | 100 | 100 | 100 |
| | | 0 < occ < 1 (%) | 0 | 0 | 0 |
| | | occ > 1 (%) | 0 | 0 | 0 |
| **Data** | **Box** | Lengths (Å) | 132.68, 135.25, 152.37 | 147.29, 142.96, 147.29 | 91.98, 121.18, 220.46 |
| | | Angles (°) | 90.00, 90.00, 90.00 | 90.00, 90.00, 90.00 | 90.00, 90.00, 90.00 |
| | **Resolution** | Supplied Resolution (Å) | 2.4 | 2.9 | 5.0 |
| | | d99 full (Masked/Unmasked) | 2.1/2.1 | 2.2/2.2 | 5.3/5.1 |
| | | d model (Masked/Unmasked) | 1.9/1.9 | 2.0/2.0 | 5.1/5.0 |
| | | d FSC model 0/0.143/0.5 (Masked) | 1.4/1.5/2.7 | 1.6/1.9/3.0 | 4.6/4.8/5.3 |
| | | d FSC model 0/0.143/0.5 (Unmasked) | 1.4/1.5/2.7 | 1.6/1.9/3.0 | 4.8/4.9/6.7 |
| | | Map min/max/mean | -0.00/1.55/0.01 | -0.00/2.19/0.01 | -0.41/0.65/0.01 |
| **Model vs. Data** | | CC (mask) | 0.86 | 0.81 | 0.72 |
| | | CC (box) | 0.85 | 0.79 | 0.73 |
| | | CC (peaks) | 0.84 | 0.78 | 0.50=9 |
| | | CC (volume) | 0.86 | 0.82 | 0.71=0 |
| | | Mean CC for ligands | 0.68 | 0.63 | |

**table S4. Library primers.**

| Name | Sequence (Partial Illumina adapter underlined) |
|---|---|
| oUni_RACE_fw | 5’ ACACTCTTTCCCTACACGACGCTCTTCCGATCTAAGCAGTGGTATCAACGCAGAGT |
| oIgG_RACE_rev | 5’ GACTGGAGTTCAGACGTGTGCTCTTCCGATCTCGAGTCGTCTTTTGGCGGGGACAGAGG |
| oIgM_RACE_rev | 5’ GACTGGAGTTCAGACGTGTGCTCTTCCGATCTGCAGGACGTTCTCACAGGAGACGAGG |
| oKappa_RACE_rev | 5’ GACTGGAGTTCAGACGTGTGCTCTTCCGATCTCCTGGGGTAGAAGCCATTCAGGAAGCAC |
| oLambda_RACE_rev | 5’ GACTGGAGTTCAGACGTGTGCTCTTCCGATCTGATGAGACACACCAGTGTGGCYTTGTTG |
| **Name** | **Sequence (overlap extension and pCHA overhangs underlined)** |
| oIGHV-1_fw | 5’ AGAAGCTTACCCATACGACGTTCCAGACTACGCTGCTAGCCAGGTMCAMCTGCTATTATT |
| oIGHV-2_fw | 5’ AGAAGCTTACCCATACGACGTTCCAGACTACGCTGCTAGCGRGGTKCAGCTGTCGGWKTC |
| oIGHV-3_fw | 5’ AGAAGCTTACCCATACGACGTTCCAGACTACGCTGCTAGCCAGGTKCAGYTGCAGGAGTC |
| oIGHV-4_fw | 5’ AGAAGCTTACCCATACGACGTTCCAGACTACGCTGCTAGCGARGTGSARYTGGTGGARTC |
| oIGHV-5_fw | 5’ AGAAGCTTACCCATACGACGTTCCAGACTACGCTGCTAGCCAGGTKCAGCTAGTMCAGTC |
| oIGHV-6_fw | 5’ AGAAGCTTACCCATACGACGTTCCAGACTACGCTGCTAGCCAGGTGCAVCTGGTDCARTC |
| oIGHV-1_rev | 5’ ctTCCACCACCTCCtgaTCCGCCGCCTCCACCGKRSGAGACGGTGACCAGGG |
| oIGHV-2_rev | 5’ ctTCCACCACCTCCtgaTCCGCCGCCTCCACCTGAGKAGACGATKRCCWGGG |
| oIGHV-3_rev | 5’ ctTCCACCACCTCCtgaTCCGCCGCCTCCACCTGAGGMGACGRTGACCAGGG |
| oIGHV-4_rev | 5’ ctTCCACCACCTCCtgaTCCGCCGCCTCCACCTRAGGMAACGGTGGKGGGRA |
| oIGKV-1_fw | 5’ CGGCGGAtcaGGAGGTGGTGGAagtGGAGGAGGTGGAtctGAYATCGTKMTKACCCARTC |
| oIGKV-2_fw | 5’ CGGCGGAtcaGGAGGTGGTGGAagtGGAGGAGGTGGAtctGAAATMATMNTTACACAKTC |
| oIGKV-3_fw | 5’ CGGCGGAtcaGGAGGTGGTGGAagtGGAGGAGGTGGAtctCAGRCYGTGGTGACYCARGA |
| oIGKV-1_rev | 5’ CTTTTGTTCTGCACGCGTGGATCCYTTGMTCTCTMGKTTGGTCC |
| oIGKV-2_rev | 5’ CTTTTGTTCTGCACGCGTGGATCCKTTRATYTCCAGATKKGTCC |
| oIGKV-3_rev | 5’ CTTTTGTTCTGCACGCGTGGATCCTTTRATYTCYAGCTYKRKYY |
| oIGKV-4_rev | 5’ CTTTTGTTCTGCACGCGTGGATCCTTTAATYTCYASYTTGGTCC |

| | |
|---|---|
| oIGKV-5_rev | 5’ CTTTTGTTCTGCACGCGTGGATCCCKTAMTYTYGMGKTTGGTCC |
| oIGLV-1_fw | 5’ CGGCGGAtcaGGAGGTGGTGGAagtGGAGGAGGTGGAtctCAGTCTRYRYTKACCCAGCC |
| oIGLV-2_fw | 5’ CGGCGGAtcaGGAGGTGGTGGAagtGGAGGAGGTGGAtctCAGCYTGYGSTGWCTCAGCC |
| oIGLV-3_fw | 5’ CGGCGGAtcaGGAGGTGGTGGAagtGGAGGAGGTGGAtctCAGWCTRYTCTGACTCARCC |
| oIGLV-4_fw | 5’ CGGCGGAtcaGGAGGTGGTGGAagtGGAGGAGGTGGAtctKCYTCTAVSCTGTCYCAGCC |
| oIGLV-5_fw | 5’ CGGCGGAtcaGGAGGTGGTGGAagtGGAGGAGGTGGAtctTCTTCTWCTGTKTCYCAGMC |
| oIGLV-6_fw | 5’ CGGCGGAtcaGGAGGTGGTGGAagtGGAGGAGGTGGAtctTCCTWTGARSTGACTCARYY |
| oIGLV-1_rev | 5’ CTTTTGTTCTGCACGCGTGGATCCKRDGACGGYCAVBYGGGTCC |
| oIGLV-2_rev | 5’ CTTTTGTTCTGCACGCGTGGATCCKGAGACGRTCAAKTGGGTHC |
| oIGLV-3_rev | 5’ CTTTTGTTCTGCACGCGTGGATCCGAGGACGGTCACCTYKKTCC |
| oIGLV-4_rev | 5’ CTTTTGTTCTGCACGCGTGGATCCTKTAACRGYCACCYGKKTCC |

| **Mastermix** | **oVH_fw** | **oVH_rev** | **oVkappa_fw** | **oVkappa_rev** | **oVlambda_fw** | **oVlambda_rev** |
|---|---|---|---|---|---|---|
| | oIGHV-1_fw | oIGHV-1_rev | oIGKV-1_fw | oIGKV-1_rev | oIGLV-1_fw | oIGLV-1_rev |
| | oIGHV-2_fw | oIGHV-2_rev | oIGKV-2_fw | oIGKV-2_rev | oIGLV-2_fw | oIGLV-2_rev |
| | oIGHV-3_fw | oIGHV-3_rev | oIGKV-3_fw | oIGKV-3_rev | oIGLV-3_fw | oIGLV-3_rev |
| | oIGHV-4_fw | oIGHV-4_rev | | oIGKV-4_rev | oIGLV-4_fw | oIGLV-4_rev |
| | oIGHV-5_fw | | | oIGKV-5_rev | oIGLV-5_fw | |
| | oIGHV-6_fw | | | | oIGLV-6_fw | |

| **Name** | **Sequence (overlap extension PCR)** |
|---|---|
| oOE_fw | 5’ AGAAGCTTACCCATACGACGTTCCAGACTAC |
| oOE_rev | 5’ CTTCTTCGGAGATAAGCTTTTGTTCTGCACGCGTGGATCC |
